# Cis-regulatory Mutations Drive Tissue-specific Subfunctionalization of sRNA loci Regulating Soybean Seed Color During Domestication

**DOI:** 10.1101/2025.01.21.634198

**Authors:** Lila O. Vodkin, Sarah I. Jones, Young B. Cho

## Abstract

Gene duplication and structural rearrangements have played a pivotal role in plant genome evolution, enabling functional divergence and the development of novel traits. In soybeans (Glycine max), the *I* locus, which regulates seed color through tissue-specific subfunctionalization of sRNA loci, offers an example of these mechanisms. Using long-read sequencing, we investigated structural variations underlying four major alleles (*I, i^i^, i^k^,* and *i*) at the *I* locus. Our study revealed that large-scale rearrangements, including duplications, inversions, and deletions, within a 180-kb CHS repeat-rich region drive allele-specific RNA silencing through small interfering RNAs (siRNAs). The dominant *I* allele contains a cis-regulatory region of DnaJ upstream of CHS genes, while the *i^i^* and *i^k^* alleles feature subtilisin- and P450-driven siRNA loci, respectively. The recessive *i* allele lacks siRNA production due to deletions or inversions disrupting CHS gene clusters. Phenotypic analyses and RNA-seq confirmed allele-specific, tissue-dependent expression of siRNAs correlating with seed coat pigmentation patterns. This study highlights the evolutionary role of repeat-rich regions in generating regulatory innovations and phenotypic diversity. Our findings underscore the importance of structural rearrangements in domestication traits and demonstrate the power of long-read sequencing for resolving complex genomic regions, advancing both evolutionary biology and crop improvement.

**SIGNIFICANCE:**

□ Naturally occurring mutations in the soybean seed coat have revealed the evolution of sRNA genes and their subfunctionalization, resulting in tissue-specific gene expression.
□ Non-homologous recombination of repetitive sequences drives cis-regulatory region rearrangements, leading to the subfunctionalization of sRNA loci that regulates one of domestication traits.

## INTRODUCTION

Gene duplication events throughout evolution have generated numerous new genes, enabling functional divergence and the emergence of novel traits in organisms. Whole-genome duplications (WGDs) are a major source of duplicated genes in flowering plants including soybeans which have undergone multiple WGDs during evolutionary history (Schmutz et al, 2010; Tang et al., 2010; Jiao et al., 2011). In addition to WGDs, smaller-scale mechanisms, such as tandem duplications (from unequal crossing over), proximal duplications (genes separated by a few others), dispersed duplications (non-adjacent and non-homologous), and retroposition (RNA-based duplication), also contribute to gene duplication (Freeling, 2009; Wang et al., 2013). Duplicated genes can follow various evolutionary paths. Most commonly, one paralog becomes a pseudogene (Lynch and Conery, 2000). Alternatively, one copy may evolve a new function (neofunctionalization), acquire a novel expression pattern (regulatory subfunctionalization), or divide ancestral functions between copies (subfunctionalization). In regulatory subfunctionalization, tissue-specific expression or regulatory responses are partitioned between duplicates, often driven by mutations in cis-regulatory regions that modify temporal, spatial, or allele-specific expression (Force et al., 1999; Gu et al., 2003). [references are not added yet on the list]

In soybeans, alleles and spontaneous mutations at the *I* locus controlling seed color, exemplify tissue-specific regulatory subfunctionalization. Early genetic studies identified four alleles—*I, i^i^, i^k^,* and *i*—at the *I* locus, arranged in a hierarchy of dominance (Bernard and Weiss, 1973; Palmer et al., 2004). Homozygous *I* plants produce yellow hilum tissue and seed coats, while *i^i^* genotypes exhibit pigmented hilum tissue on a yellow seed coat. The rare *i^k^* allele extends pigmentation beyond the hilum tissue, creating a saddle-shaped pattern, and homozygous *i* plants have fully pigmented hilum and seed coat tissues (see Figure 1). Domestication favored soybeans with less pigmented seeds with *I* and *i^i^* alleles (Li et al., 2013; Zhou et al., 2015; Valliyodan et al., 2016).

**Figure 1.**
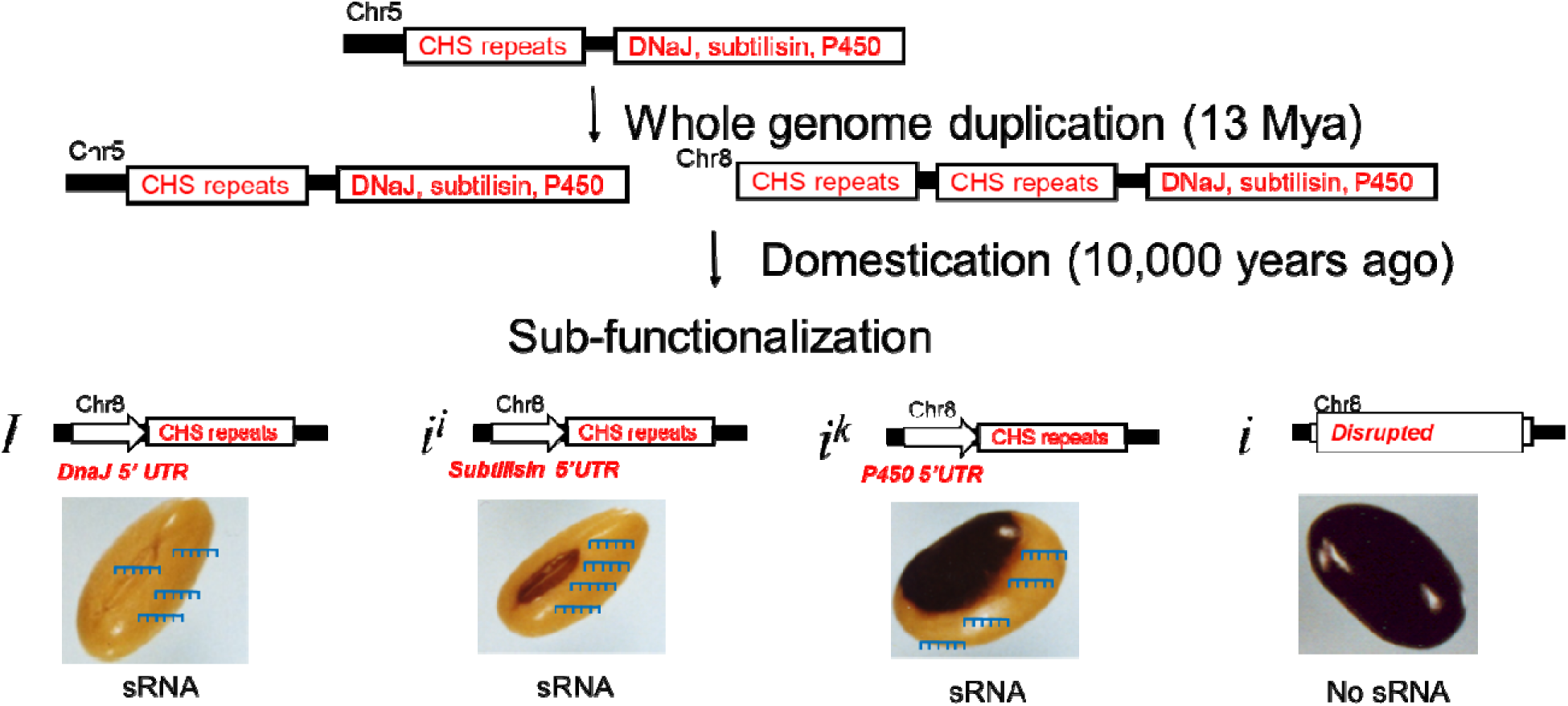
Graphical representation of sRNA locus evolution regulating soybean seed color, drives cis-regulatory mutations, leading to tissue-specific subfunctionalization of the four I alleles.

While the exact mechanism of tissue-specific subfunctionalization at the *I* locus remains unclear, significant progress has been made in understanding seed color regulation. The *I* locus produces siRNAs that downregulate chalcone synthase (CHS) mRNAs, a key enzyme for flavonoid and pigment production (Wang et al., 1994; Todd and Vodkin, 1996; Tuteja et al., 2004, 2009; Senda et al., 2004, 2012; Cho et al., 2013, 2017; Jia et al., 2020). Additionally, structural changes in the *I* region, such as gene copy number variations, contribute to this regulation (Clough et al., 2004; Tuteja et al., 2008; Cho et al., 2019; Xie et al., 2019; Liu et al., 2020). BAC sequence analysis revealed that the I locus in Williams 82 contains 12 functional CHS genes within a 180-kilobase region, many of which are arranged in 10-kb inverted repeats (Clough et al., 2004). A 5’ partial subtilisin gene fragment near one of the CHS repeat clusters has been implicated in transcribing CHS siRNA precursors (Clough et al., 2004), with recent studies confirming its necessity for CHS siRNA production, particularly in black-seeded wild soybeans (Xie et al., 2019; Liu et al., 2020).

The repetitive nature of CHS genes at the *I* locus presents a challenge for genome assembly using short-read sequencing methods. The first two soybean genome assemblies lacked a complete *I* locus (Schmutz et al, 2010; see Wm82.a1 and Wm82.a2 in soybase.org), necessitating the use of BAC sequences to fill in gaps (Cho et al., 2019). Short-read alignments identified NAHR between distant CHS genes as the cause of ii to i mutations, deleting 138 kb of sequence and reducing the number of CHS genes from 12 to a single hybrid gene (Cho et al., 2019). The recent genome assembly incorporates long-read sequencing to provide a more comprehensive reference (Wm82.a4, soybase.org). This version includes a new gene model, 08G110380, which encodes a chimeric gene transcribing a 5’ partial subtilisin gene fragment, CHS1, and CHS3, further supporting the role of structural rearrangements in driving tissue-specific siRNA production (Clough et al., 2004; Xie et al., 2019; Liu et al., 2020).

In this study, we used long-read sequencing (Pacific Biosciences) to understand the evolution of the *I* locus across multiple lines with different alleles. Our results revealed distinct origins of RNA silencing mechanisms, all driven by structural variations, including inversions, duplications, and deletions within the 180-kb region containing up to 12 CHS genes. These structural rearrangements generated various non-*CHS* gene fragments (e.g., subtilisin, DnaJ, or P450) with intact promoters upstream of *CHS* inverted repeats, resulting in the four major alleles (*I, i^i^, i^k^,* and *i*) (Summarized in Figure 1). This study underscores the role of structural rearrangements in repetitive regions in driving phenotypic diversity at the *I* locus, supporting the hypothesis that gene duplication and repetitive regions facilitate cis-regulatory mutations leading to tissue-specific subfunctionalization of small regulatory RNA (sRNA) loci, changing the domestication trait.

## RESULTS

### PCR Screening Reveals Large NAHR Events Affecting the Chimeric sRNA-Generating Gene (08G110380)

Large non-allelic homologous recombination (NAHR) events deleted 138 kb region including the chimeric sRNA-generating gene (08G110380) in three spontaneous black mutations (*i* allele) (Cho et al., 2019). To further distinguish these large NAHR events from smaller mutations, we conducted PCR screening of 30 seed color mutants from the soybean germplasm collection (Supplemental Data Set 1), revealed that large NAHR events accounted for 52% of the *i^i^* to *i* mutations (Table 1). The large NAHR event between the two most distal CHS genes forms a hybrid CHS5:1 gene, detectable by a 7.9 kb amplicon in mutant lines but absent in the parental Williams cultivar (Supplemental Figure 1). PCR assays confirmed the presence of hypothetical (Glyma.08G110600) and RRM (Glyma.08G109600) gene amplicons in the yellow seed black hilum parental lines (W43 and UC7), but these were absent in some black seed mutants (W55, W130 and UC9), where the 7.9-kb NAHR amplicon was instead present (Supplemental Figure 1 and 2). However, the remaining 16 black mutants retained the hypothetical and RRM gene amplicons but lacked the 7.9-kb band, suggesting smaller mutations (Table 1). This screening allows us to focus on lines with smaller mutations that may affect the chimeric sRNA-generating gene (08G110380) in distinct ways.

**Figure 2.**
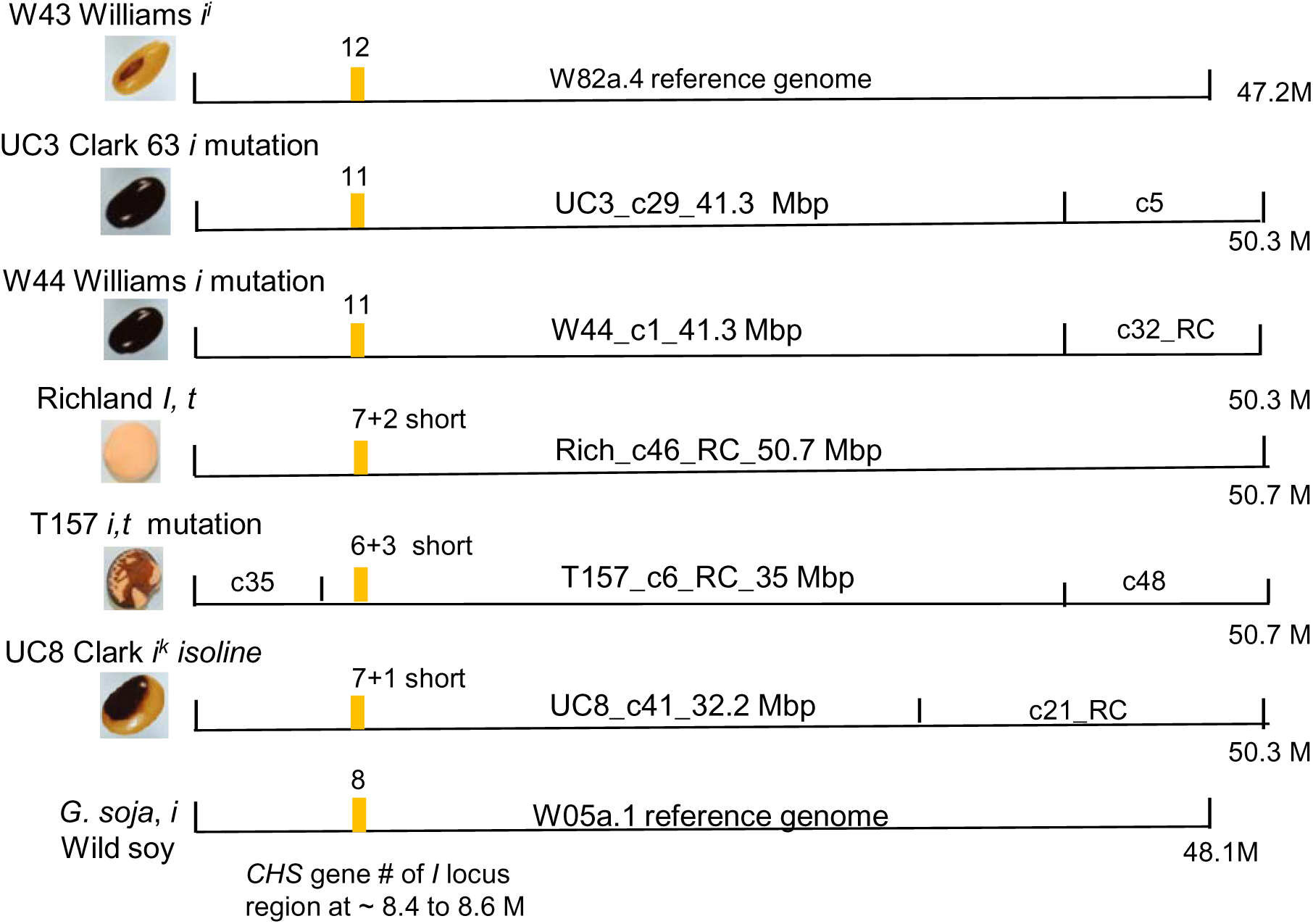
Diagram of Contig Sizes and the Number of *CHS* genes on Chromosome 8 in each Assembly for the Different *I* Alleles or Mutations Examined. The exact lengths of contigs in millions of bp (M) are shown; c, contig number; RC reverse complement. BLAST results are shown in Supplemental File 2. The number above the yellow bar is the number of *CHS* genes located in the *I* locus region of each Chromosome 8 as determined in Supplemental Table 1 and Supplemental Data Set 3. Short refers to truncated *CHS* copies of various sizes. Both Richland and T157 also carry the recessive *t* allele. T157 is a spontaneous mutation from Richland. The homozygous recessive *t* allele is a flavonoid 3’ hydroxylase (Zabala and Vodkin 2003) that also reduces the intensity of the pigment in the seed coat and further has an epistatic interaction that produces cracks in the pigmented seed coat and exposes the cotyledons beneath as shown in the example seed picture.

**Table 1.** PCR characterization of independent color mutations.

| Mutant Lab No. | Year Found | Mutant Color | Hypo Gene | RRM Gene | 7.9 kb PCR | NAHR Pattern |
| --- | --- | --- | --- | --- | --- | --- |
| *(a) Seed color mutations consistent with 138-kb deletion events* |  |  |  |  |  |  |
| W55 | 1975 | *i*, black | No | No | Yes | Yes |
| W130 | 1980 | *i*, black | No | No | Yes | Yes |
| UC9 | 1965 | *i*, black | No | No | Yes | Yes |
| UC801 | 1971 | *i*, black | No | No | Yes | Yes |
| UC803 | 1970 | *i*, black | No | No | Yes | Yes |
| UC805 | 1955 | *i*, black | No | No | Yes | Yes |
| UC810 | 1956 | *i*, black | No | No | Yes | Yes |
| UC811 | 1985 | *i*, black | No | No | Yes | Yes |
| UC814 | 1947 | *i*, black | No | No | Yes | Yes |
| UC815 | 1954 | *i*, black | No | No | Yes | Yes |
| UC818 | 1989 | *i*, black | No | No | Yes | Yes |
| UC819 | 1959 | *i*, black | No | No | Yes | Yes |
| UC820 | 1985 | *i*, black | No | No | Yes | Yes |
| UC821 | 1980 | *i*, black | No | No | Yes | Yes |
| *(b) Seed color mutations not consistent with 138-kb deletions* |  |  |  |  |  |  |
| UC804 | 1953 | *i*, brown | Yes | Yes | No | No |
| UC806 | 1975 | *i*, black | Yes | Yes | No | No |
| UC807 | 1947 | *i*, brown | Yes | Yes | No | No |
| UC808 | 1986 | *i*, black | Yes | Yes | No | No |
| UC809 | 1954 | *i*, black | Yes | Yes | No | No |
| UC812 | 1966 | *i*, black | Yes | Yes | No | No |
| UC813 | 1969 | *i*, brown | Yes | Yes | No | No |
| UC816 | 1954 | *i*, black | Yes | Yes | No | No |
| UC822 | 1966 | *i*, black | Yes | Yes | No | No |
| UC823 | 1966 | *i*, black | Yes | Yes | No | No |
| UC825 | 1971 | *i*, black | Yes | Yes | No | No |
| UC3* | 1954 | *i*, black | Yes | Yes | No | No |
| W44* | 1980 | *i*, black | Yes | Yes | No | No |
| T157*<sup>a</sup> | 1938 | *i*, im black | Yes | Yes | No | No |
| UC817 | 1961 | *i*, buff | Yes | Yes | No | No |
| UC824 | 1976 | *i*, im black | Yes | Yes | No | No |

\* indicates three lines also sequenced by Pac Bio in this report; aT157 determined by sequencing instead of PCR. im black, imperfect black. The Hypo gene is Glyma.08G11060, and the RRM gene is Glyma.08G109600 in the Wm82a.4 genome. Supplemental Data Set 1 has the searchable accession numbers and full phenotypic information for the parent and mutant cultivars.

### Short Read Sequencing Does Not Reveal Smaller Mutations, Necessitating the Use of Long Read Technologies for the Repetitive I Locus Regions

While large deletions from NAHR events were easily detected by short-read Illumina data aligned to the Williams 82 reference genome (Cho et al., 2019), this was not the case for other mutant lines, such as Williams 44 (W44, *i* allele), which did not show PCR patterns consistent with large NAHR events (Table 1). CHS sRNAs were absent in W44, yet short-read re-sequencing failed to reveal any structural variations affecting the chimeric sRNA-generating gene (08G110380) (Supplemental Figure 3). To address the limitations of short-read sequencing in detecting smaller mutations, we used long-read PacBio sequencing on *i^i^* to *i* mutations (UC3 and W44), the Richland *I* parent line and its T157 mutation, and UC8, an isoline with the *i^k^* saddle pattern allele. Sequencing from two flow cells per variety yielded 8.9 to 41.8 Gb with raw read lengths of 10.3 to 18.0 kb (Table 2; Supplemental Data Set 2A). Assembled genome sizes ranged from 1.020 to 1.038 Gb, with N50 contig sizes between 31.6 and 43.5 Mb and coverage depths of 41x to 77x, indicating high-quality assemblies (Table 2; Supplemental Data Set 2B).

**Table 2.**
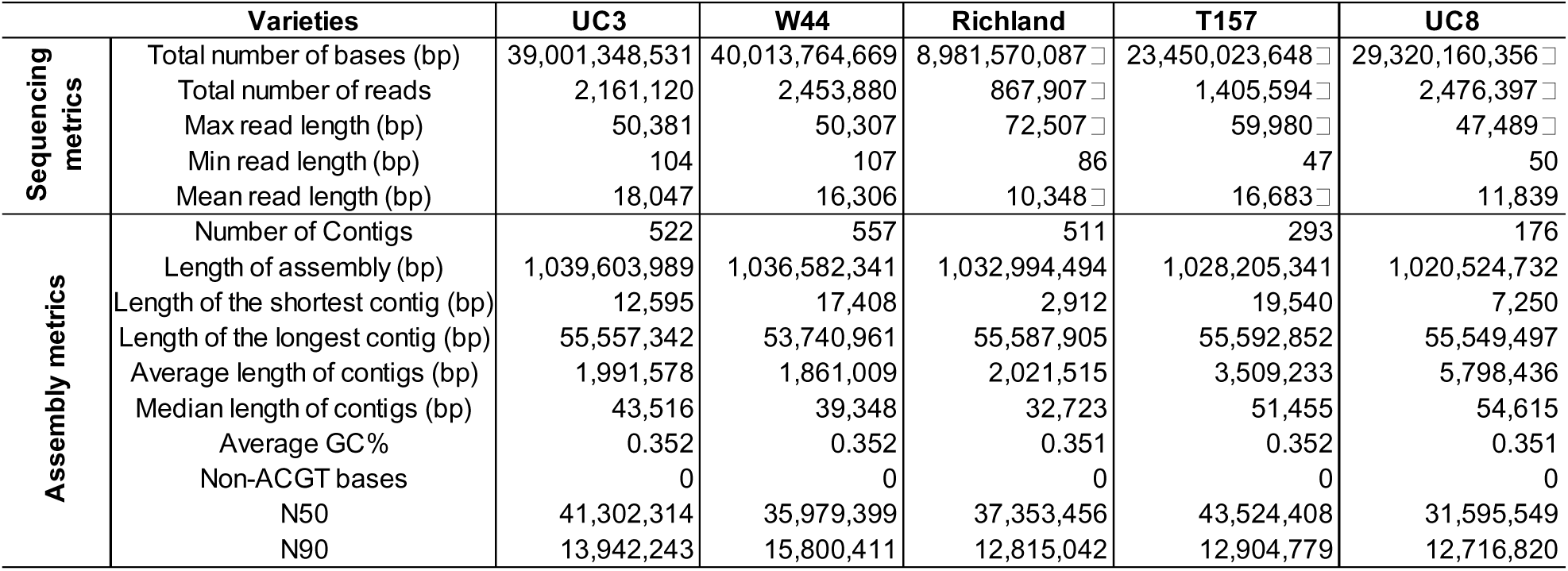
Summary of PacBio sequencing and genome assembly statistics for soybean varieties.

| Varieties |  | UC3 | W44 | Richland | T157 | UC8 |
| --- | --- | --- | --- | --- | --- | --- |
| Sequencing metrics | Total number of bases (bp) | 39,001,348,531 | 40,013,764,669 | 8,981,570,087 □ | 23,450,023,648 □ | 29,320,160,356 □ |
|  | Total number of reads | 2,161,120 | 2,453,880 | 867,907 □ | 1,405,594 □ | 2,476,397 □ |
|  | Max read length (bp) | 50,381 | 50,307 | 72,507 □ | 59,980 □ | 47,489 □ |
|  | Min read length (bp) | 104 | 107 | 86 | 47 | 50 |
|  | Mean read length (bp) | 18,047 | 16,306 | 10,348 □ | 16,683 □ | 11,839 |
| Assembly metrics | Number of Contigs | 522 | 557 | 511 | 293 | 176 |
|  | Length of assembly (bp) | 1,039,603,989 | 1,036,582,341 | 1,032,994,494 | 1,028,205,341 | 1,020,524,732 |
|  | Length of the shortest contig (bp) | 12,595 | 17,408 | 2,912 | 19,540 | 7,250 |
|  | Length of the longest contig (bp) | 55,557,342 | 53,740,961 | 55,587,905 | 55,592,852 | 55,549,497 |
|  | Average length of contigs (bp) | 1,991,578 | 1,861,009 | 2,021,515 | 3,509,233 | 5,798,436 |
|  | Median length of contigs (bp) | 43,516 | 39,348 | 32,723 | 51,455 | 54,615 |
|  | Average GC% | 0.352 | 0.352 | 0.351 | 0.352 | 0.351 |
|  | Non-ACGT bases | 0 | 0 | 0 | 0 | 0 |
|  | N50 | 41,302,314 | 35,979,399 | 37,353,456 | 43,524,408 | 31,595,549 |
|  | N90 | 13,942,243 | 15,800,411 | 12,815,042 | 12,904,779 | 12,716,820 |

The phenotypes and CHS gene counts (8 to 12) on chromosome 8 across the five assembled genomes (Figure 2; Supplemental Table 1). Local BLAST searches identified the contig with the highest homology to CHS4, exclusively on chromosome 8 (Supplemental Table 1; Supplemental Data Set 3). Chromosome 8 sizes in the five genomes ranged from 50.30 to 50.65 Mbp, similar to the Wm82a.4 reference genome’s 47.2 Mbp but without gaps or Ns. W44 and Richland show full-length chromosome 8, while the other three lines have the *I* locus region within a single contig containing 8 to 12 CHS genes (Figure 2; Supplemental Table 1). The SHMT gene (serine hydroxymethyl transferase, Glyma.08G108900.3) is positioned between 8.39 and 8.43 Mbp in all genomes, comparable to 8.36 Mbp in Wm84a.4. The SHMT-to-galactosidase region spans 178,466 to 193,112 bp, close to the 186,027 bp in Wm82a.4, manually annotated using BLAST (Supplemental Data Set 4).

### Long-Read Sequencing Reveals that Deletion of the CHS1 Region from the Chimeric sRNA-Generating Gene (08G110380) in the UC3 Black Mutation (*i* allele)

PacBio long-read sequencing compares the assembly sequences of two mutant lines: one with a dominant yellow seed coat and black hilum tissue phenotype (*i^i^*) and another with a recessive black seed coat and hilum tissue (*i*). UC3 (PI 547438), a self-black mutation identified in the inbred variety Clark 63 in 1954, has an identical SHMT-to-Gal region to the Williams reference genome, except for a 5,498 bp deletion encompassing the CHS1 gene, part of the CHS1,3,4 inverted repeat cluster located just after a partial subtilisin fragment (Figure 3A). To confirm this was not an assembly error, we conducted a BLAST search using over 2 million raw reads, with an average length of 18,047 bp. Several reads (18,237, 21,693, and 16,153 bp) confirmed the exact deletion within the larger sequence (Figure 3B; Supplemental Data Set 5). Further support for the assembly’s accuracy came from HindIII restriction mapping, which showed a 9.5 kb fragment in the UC3 PacBio assembly, matching previous RFLP analysis that identified a 9.4 kb HindIII fragment lacking the CHS1 promoter in UC3 (Todd and Vodkin, 1996). The deletion of the antisense-oriented CHS1 gene is likely crucial for the chimeric gene (08G110380), as it may disrupt the long-inverted repeat responsible for producing small RNAs in the yellow seed coat with black hilum phenotype (*i^i^*).

**Figure 3.**
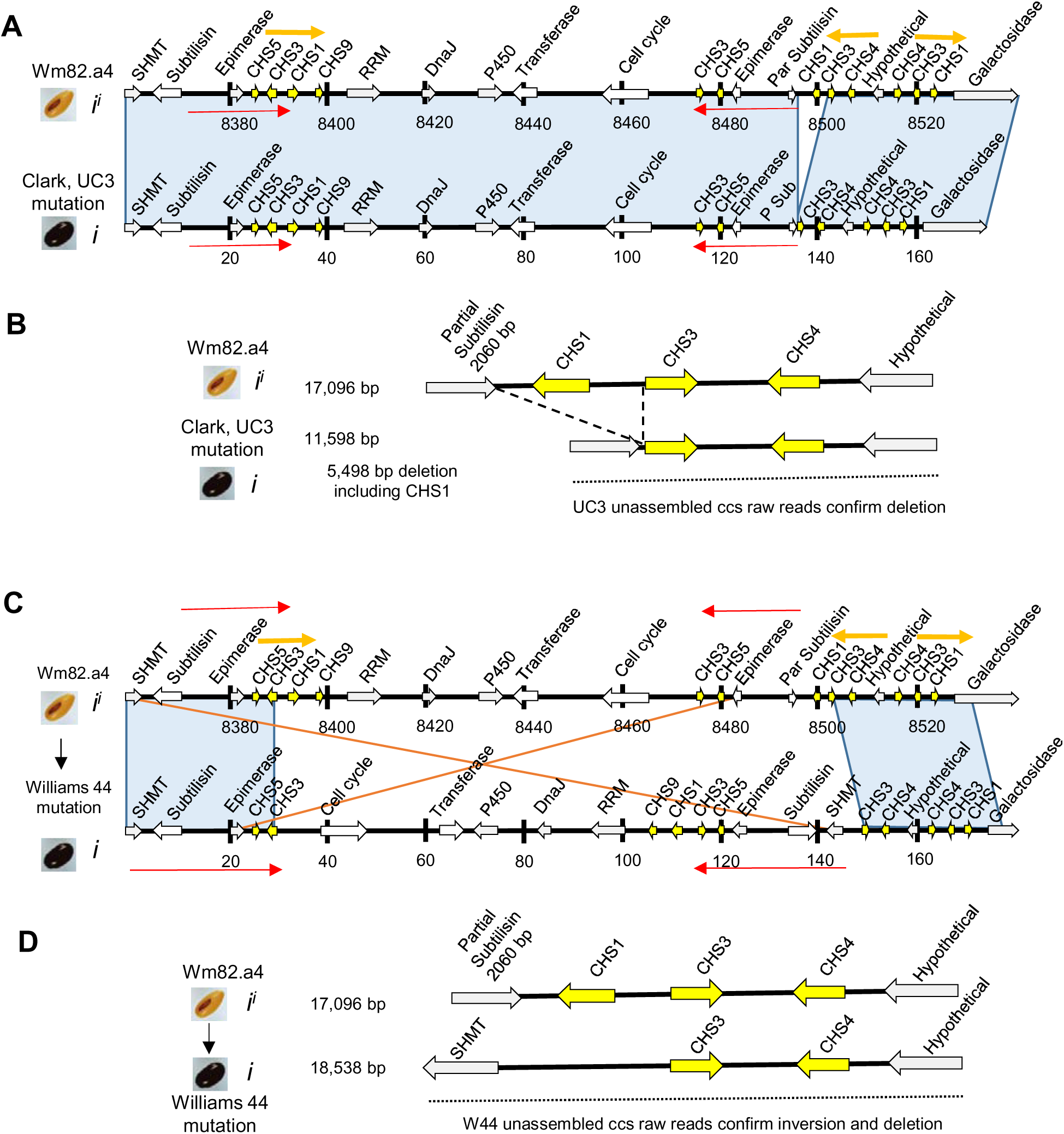
Assembly of Two *i^i^* to *i* Mutations that Show Deletion or Inversion Arrangements. Graphical representations of the positions of the gene models were determined by BLAST for the segments from the SHMT to galactosidase gene models as shown in Supplemental Data Set 4. Numbering is anchored at the beginning of the SHMT gene model for each graph in basepairs. Blue shading indicates similar gene orders. Red and orange arrows indicate large, duplicated regions which involve *CHS* or nearby genes. **(A)** Comparison of the Wm82a.4 reference genome with the assembled UC3 (*i^i^* to *i* black mutation) genome. **(B)** Enlargement of the region surrounding the 5,498 bp deletion region found in UC3. Horizontal dotted lines represent the area included within larger UC3 raw reads as shown in Supplemental Data Set 5 that confirm the deletion area as shown. **(C)** Comparison of the Wm82a.4 reference genome with the W44 mutation which is a spontaneous isogenic *i^i^* to *i* black mutation in the Williams (W43) parent. Red arrows in W44 show the inverted region. **(D)** Enlargement of the region surrounding the inversion area that removes the partial subtilisin and the antisense *CHS1* gene copy of the *CHS1,3,4* repeat cluster. Horizontal dotted lines represent the area included within raw reads from the W44 PacBio data as shown in Supplemental Data Set 6, which confirms the inversion order as shown.

### A Large Inversion Removed the Partial Subtilisin Fragment and CHS1 from the Chimeric Gene (08G110380) in the Williams 44 Black Mutation (*i* allele)

The Williams 44 (W44, *i* allele) mutation resulted from a large inversion that rearranged several gene models (Figure 3C and D). This inversion removed both the partial subtilisin fragment and the CHS1 gene from the chimeric gene (08G110380). Two long reads, each over 21,000 bp, revealed that the duplicated SHMT gene is now located near CHS3 and CHS4 (Figure 3D; Supplemental Data Set 6). Not detectable through short-read sequencing alone (Supplemental Figure 3), this inversion highlights the essential role of the CHS1 gene and the partial subtilisin fragment in the chimeric gene (08G110380) for small RNA production.

### The Dominant *I* Allele Contains a Different Chimeric Gene with a Partial DnaJ Fragment Adjacent to CHS Genes, While the *I* to *i* Mutation Truncates the CHS Genes

The structure of the dominant *I* allele in Richland, which produces a yellow seed coat and hilum, has a gene order similar to the recently published W05 reference genome (wild, small-seeded black variety) rather than the Williams 82 reference genome (*i^i^* allele, Wm82a.4). Richland contains nine CHS gene copies, compared to 12 in Wm82a.4, with an inverted region that includes an extra RRM copy and a partial DnaJ gene adjacent to three closely spaced, truncated CHS genes (Figure 4A; Supplemental Table 3). The only structural difference between the PacBio sequence of T157 (isogenic to Richland, with a spontaneous *I* to *i* mutation) and Richland lies in the CHS genes near the DnaJ fragment. In both Richland and T157, the DnaJ fragment is 406 bp and abuts the CHS genes (Figure 4B). In Richland, it is followed by two truncated CHS genes in inverted orientation, separated by 148 bp, and a full-length CHS1 gene. The T157 mutation features a 2,454 bp deletion, further truncating the first and third CHS genes to 251 bp and 330 bp, respectively, and eliminating an intergenic region. This deletion likely disrupts dsRNA precursor formation. Interestingly, the orientation of the first CHS gene in the *I* allele is in the plus orientation, unlike the antisense CHS1 gene in ii alleles. The regions from the DnaJ fragment to the start of the galactosidase gene in both Richland and T157 were confirmed in raw reads with the same gene order (Figure 4B; Supplemental Table 3; Supplemental Data Sets 7 and 8). HindIII restriction mapping shows that a 12.6 kb fragment in Richland was replaced by a 10.1 kb fragment in T157, consistent with previous RFLP analysis (Todd and Vodkin, 1996).

**Figure 4.**
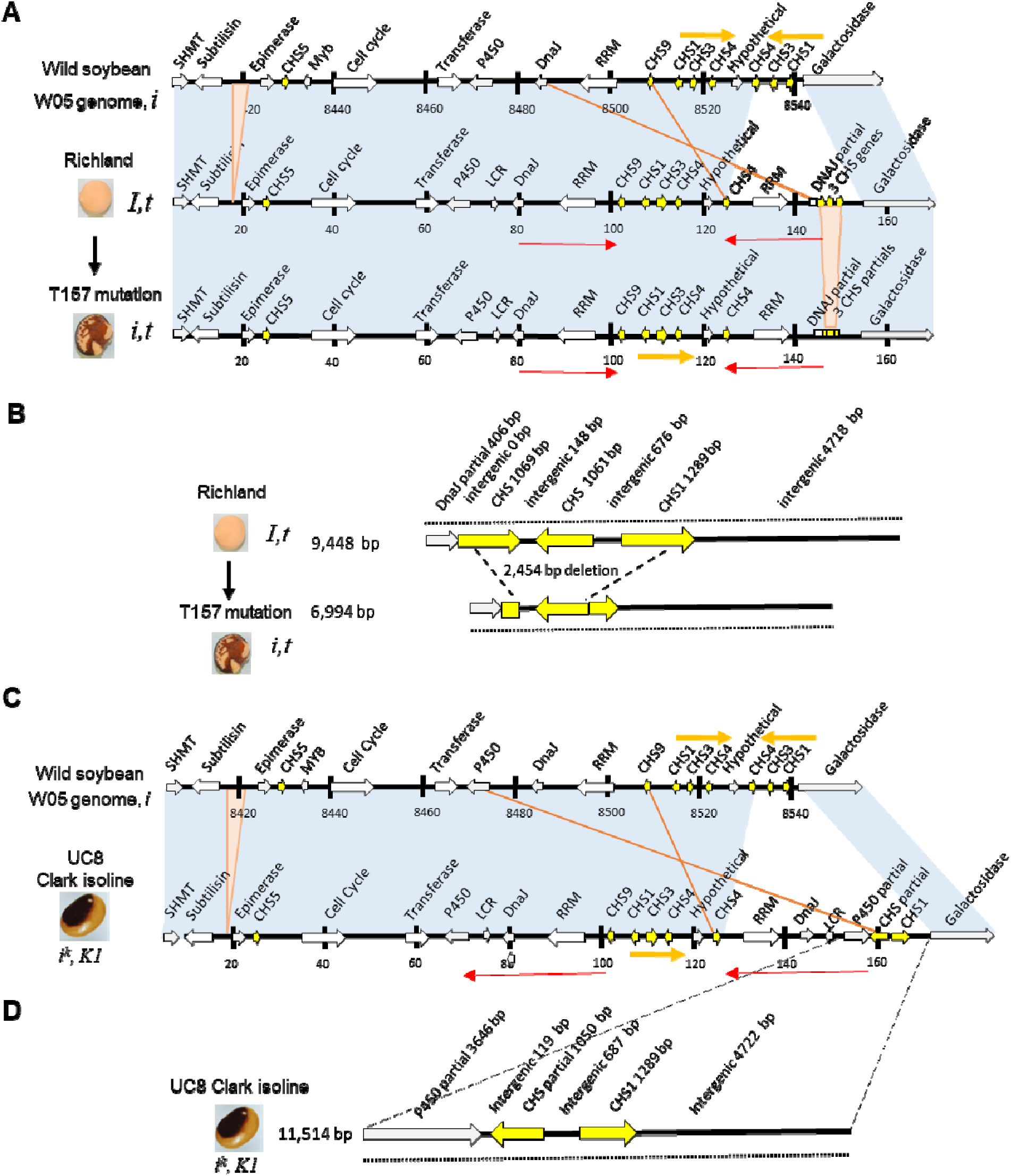
Assemblies of *I* and *i^k^* Alleles Reveal Two Different Origins of Small RNA Generation. Graphical representations of the positions of the gene models were determined by BLAST for the segments from the SHMT to galactosidase gene models as shown in Supplemental Data Set 4. Numbering is anchored at the beginning of the SHMT gene model in basepairs. Blue shading indicates similar gene orders. Pink and white triangles are major structural differences. Red and orange arrows indicate large, duplicated regions which involve *CHS* or nearby genes. **(A)** Comparison of the W05 wild soybean reference genome (Xie, et al. (2019) to the PacBio assembled contig for Richland (*I*, *t* yellow hilum and seed coat) and of the Richland mutation T157 (*i, t*, with imperfect black and defective seed coats). The small region highlighted in pink denotes the only major structural difference between Richland and T157. **(B)** Enlargement of the region surrounding the 2,454 bp deletion region found in T157. Horizontal dotted lines represent the area included within larger T157 raw reads as shown in Supplemental Data Sets 7 and 8 that confirm the arrangements of these genes in Richland and T157. **(C)** Comparison of the W05 wild soybean reference genome to the PacBio assembled contig for UC8, a Clark isoline containing the *i^k^* allele that specifies a black saddle phenotype as shown. **(D)** Enlargement of the region encompassing the region from the partial P450 gene that now abuts two *CHS* genes in inverted orientations. The horizontal dotted line indicates confirmation of this arrangement in raw reads as shown in Supplemental Data Set 9.

### The *i^k^* Allele Contains a Third Chimeric Gene with a Partial P450 Fragment, Defining the Saddle Pattern Specific sRNA Expression

UC8 (PI 547450), a black saddle isoline released in 1970, was developed by backcrossing the *i^k^* allele from the non-adapted Black Eyebrow variety into the recurrent parent Clark 63 (*i^i^*, yellow seed coat with black hilum) for six generations (GRIN). Similar to the Richland and T157 genomes, UC8 gene order from SHMT through CHS4 in the CHS4,3,1 cluster aligns more closely with the wild soybean W05 than with Clark 63 (*i^i^* allele) (Figure 4C). Due to phenotype-based selection, backcrossing can introduce large regions from the donor genome. Interestingly, the *i^k^* allele exhibits a third structural arrangement for sRNA generation, distinct from Richland and T157 (Figure 4C). This region includes an inverted duplication of several genes, such as the RRM gene, a full DnaJ gene, and a partial P450 gene, followed by two inverted CHS genes in a head-to-head orientation, unlike the tail-to-tail orientation in Richland. The two main differences between Richland and UC8 are the P450 driver and the absence of the first partial CHS gene in UC8 (Figure 4D). The 11,514 bp region containing this novel P450-CHS arrangement was confirmed by larger raw reads in UC8 (Supplemental Data Set 9).

### Upstream Promoter Regions of CHS siRNA-Generating Regions and Their Cognate Genes are Nearly Identical

The genomic structures of the three alleles reveal different potential drivers of CHS siRNA expression. To assess mutations in the promoter regions, we compared the upstream sequences of the functionally complete (cognate) genes with those of the duplicated regions driving precursor siRNA expression. In the *I* allele, the 8,817 bp region upstream of the partial DnaJ fragment in Richland was identical to that of the cognate *DnaJ* gene (*Glyma.08G109700*). Similarly, the 8,776 bp region upstream of the partial *P450* fragment in the *i^k^* allele matched the cognate *P450* gene (*Glyma.08G109900*). These sequences are subsets of the duplicated regions (Figure 4, red arrows), suggesting similar promoter strength unless larger chromatin context is involved. In UC3 (*i^i^* to *i* mutation), the 9,809 bp intergenic region between the partial *subtilisin* gene and the flanking *epimerase* gene is identical to the cognate gene (*Glyma.08G109000*), except for four ATAT bases in an upstream AT-rich region. The W44 genome (*i^i^* to *i* mutation) also shows two identical 9,809 bp regions between the two *subtilisin* copies (Supplemental Figure 4).

### Differential Expression of Subtilisin (Glyma.08G109000) and Chimeric Gene (Glyma.08G110380) Correlates with the ii Allele Phenotype: Yellow Seed Coat with Black Hilum

Long-read assemblies identify three CHS siRNA-generating regions, suggesting a significant role for flanking subtilisin, DnaJ, or P450 fragments adjacent to inverted repeat CHS genes. Promoters must account for the allele-specific expression patterns of CHS siRNAs in *i^i^* (subtilisin-driven) and *i^k^* (P450-driven) alleles, as well as the dominance hierarchy: *I* (DnaJ-driven) > *i^i^* (subtilisin-driven) > *i^k^* (P450-driven) > *i* (fully pigmented, no CHS siRNAs). RNA-seq analysis of hila and seed coats revealed differential expression in the subtilisin-driven ii allele (Figure 5; Supplemental Table 4; Supplemental Data Set 10). The chimeric gene (Glyma.08G110380, 10,340 bp) spans coding and intergenic regions from the partial subtilisin to the antisense CHS1 gene and through CHS3 in yellow seed coats (Figure 5A), with lower expression in the hila. This partial subtilisin gene corresponds to the first 2061 bases and includes 4 of the 11 exons of the cognate subtilisin gene (Supplemental Figure 5). The final 378 bases align with an intergenic region between CHS3 and CHS4. Chimeric reads between the partial subtilisin gene and the intergenic region suggest these are read-through transcripts (Figure 5C, D). The cognate subtilisin gene (Glyma.08G109000) is expressed 3.2-fold higher in non-pigmented seed coats compared to pigmented hila (25.2 vs 7.8 RPKM, Figure 5B). Similarly, P450 gene (Glyma.08G109900) expression is higher in non-pigmented seed coats, while SHMT expression is elevated in pigmented hila. Other genes near the I locus, including DnaJ (Glyma.08G109700), show no significant expression differences between tissues. These results suggest that differential expression of the cognate subtilisin gene, and possibly the partial subtilisin gene, correlates with the *i^i^* allele’s seed coat pigmentation pattern.

**Figure 5.**
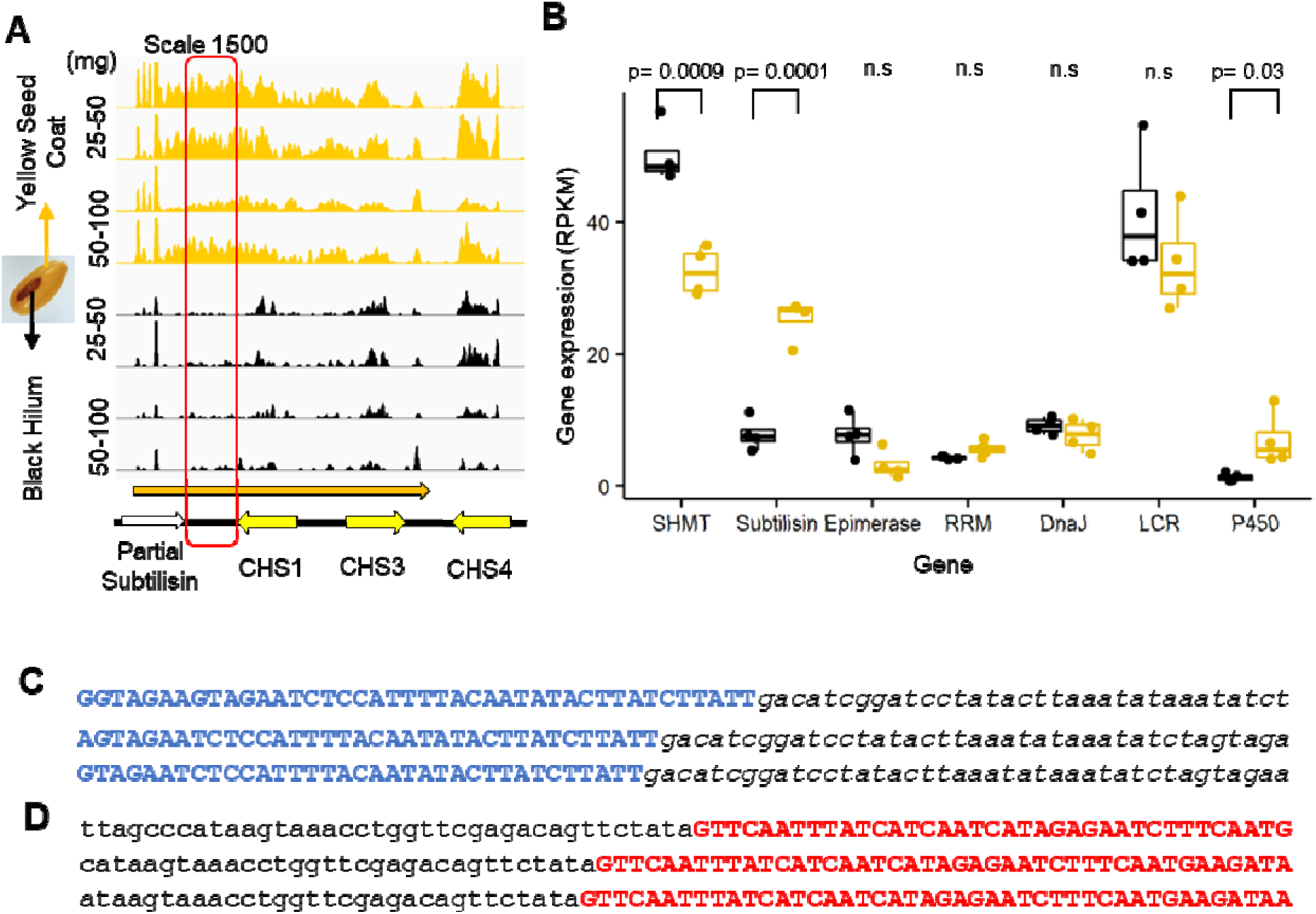
Differential expression of *CHS* siRNA precursors and the cognate subtilisin gene within pigmented hila versus the yellow seed coats of the *i^i^* pattern genotype. Supplemental Data Set 10 has the complete tissue stages and all other information including raw counts and normalized reads. **(A)** Alignments showing normalized read counts viewed with the IGV browser of independent RNA Seq data sets (R number) from Williams seed coats minus hilum (yellow color) or from the dissected hila (black color) from the indicated immature seed weight ranges. The read counts were normalized by multiplying each value by [1,000,000 / (total Read Count)]. Alignments were made to the full SHMT to Gal region (182,027 bp) but only the *CHS* siRNA precursor region are shown. A long transcript (orange arrow) from the 5’ partial subtilisin through *CHS3* (Glyma.08G110380, 10340 bp) is more highly expressed in the yellow seed coat samples than in the black hilum. The red box represents the unique intergenic region between the partial subtilisin and *CHS1* (Glyma.08G110420). **(B)** Gene expression comparisons of RNA-seq for genes within the *I* locus region. SHMT (p=0.0009; 1.5 fold; 50.2 vs 32.6), subtilisin (p=0.0001; 3.2 fold; 7.8 vs 25.2) and P450 (p=0.03; 3 fold; 1.3 vs 6.9) are significantly different between the hila (black plots) versus the seed coat proper (yellow plots). The box plots show the median (central line), the lower and upper quartiles (box), and the minimum and maximum values (whiskers). Each dot represents a value (n=4). n.s stands for non-significantly different when p>0.05. ANOVA is used for statistical analysis. The four repeats for statistical analysis combined the two stages of 25-50 mg and 50-100 mg immature seed weight ranges as shown in part (A) above and thus represent tissue between the 25-100 mg weight range. **(C)** Three example RNA-seq reads that illustrate chimeric sequences consisting of the end of the partial subtilisin gene region (bold blue font) and the intergenic region (small letters in italics font). Additional junction sequences are shown in Supplemental File 6. **(D)** Three example RNA-seq reads that illustrate chimeric sequences consisting of the intergenic region (small letters in italics font) that is contiguous with the *CHS1* antisense gene (red bold font). Additional junction sequences are shown in Supplemental File 6.

## DISCUSSION

The soybean genome has undergone two rounds of whole-genome duplication: one approximately 59 million years ago (Mya), shared with legumes like Medicago and Lotus, and a second, lineage-specific duplication around 13 Mya (Schlueter et al., 2004; Schmutz et al., 2010; Vanneste et al., 2014; Liu et al., 2015). These duplications have shaped the soybean genome, resulting in ∼75% of its genes having multiple paralogs (Schlueter et al., 2007; Schmutz et al., 2010; Severin et al., 2011; Singh and Jain, 2015; Han et al., 2016; Du et al., 2023; Duan et al., 2023). The more recent duplication relocated a CHS gene repeat from chromosome 5 to chromosome 8, forming a 180-kb CHS repeat-rich region on chromosome 8 (Supplemental Figure 6, Schmutz et al, 2010). This study demonstrated that the CHS repeat region has driven further genomic rearrangements, including promoter modifications and tissue-specific CHS siRNA expression (Figures 2–6). A genomic inversion during domestication brought a partial subtilisin fragment near the CHS1,3,4 cluster, forming the chimeric gene 08G110380 and leading to the yellow seed color of *i^i^* allele (Figures 3–4), which persists in modern cultivars (Liu et al., 2020). It is unclear why domestication favored soybeans with yellow seed (Li et al., 2013; Zhou et al., 2015; Valliyodan et al., 2016). We speculate that ancient farmers prefer lower flavonoid and anthocyanin concentrations in seeds (Todd and Vodkin, 1993), giving a less bitter taste and better digestibility (Žilić et al., 2013; Zhang et al., 2022).

The domesticated yellow soybean (*i^i^* allele) can revert to a black seed coat (*i* allele), a pre-domestication trait involving the CHS repeat-rich region. We analyzed 30 independent *i^i^* (yellow seed coat with black hilum) to *i* (fully pigmented seed coat) mutations from the soybean germplasm collection using PCR (Table 1; Supplemental Figures 1 and 2). Fourteen of these mutations were linked to large 138-kb deletions by non-allelic homologous recombination (NAHR), abolishing the chimeric gene (08G110380). For smaller mutations, long-read sequencing provided detailed structural insights (Figure 2). In the UC3 mutation, the deletion of the antisense CHS1 sequence of the chimeric gene (08G110380) pinpointed this region as the source of small RNAs (Figure 3A and B). Another mutation (Wm44) involved a large inversion that removed both the partial subtilisin and antisense CHS1 sequence of the chimeric gene (08G110380) and duplicated copies of the SHMT and subtilisin genes (Figure 3C and D). This inversion separates the partial subtilisin-*CHS1* region from the *CHS1,3,4*, cluster. The arrangement of large 10-20 kb regions of identical or nearly identical sequences (shown with red and yellow arrows in Supplemental Figure 1) containing multiple *CHS* copies, as well as duplication of the epimerase and part of the subtilisin genes, can spawn both large inversions as well as the large frequent 138-kb NAHR deletions, that led to loss of the entire repeat region.

Genome rearrangements around the CHS repeat regions drive tissue-specific expression patterns of CHS siRNAs in the *i^i^* (subtilisin-driven) allele and establish the dominance hierarchy: *I* (DnaJ-driven) > *i^i^* (subtilisin-driven) > *i^k^* (P450-driven) > *i* (fully pigmented, no CHS siRNAs). In the *i^i^* allele, the significantly lower expression of the subtilisin gene and CHS siRNA precursors in the pigmented hilum compared to the yellow seed coat correlates with the tissue-specific pigmentation phenotype (Figure 5). This suggests that CHS siRNA expression in the *i^i^* allele is governed by the partial expression of subtilisin. In the *I* allele (yellow hilum and seed coat), CHS siRNAs are driven by a truncated DnaJ fragment instead of the partial subtilisin gene. This fragment is positioned upstream of two slightly truncated inverted CHS genes (Figure 4A, B). A similar structure was identified in the Japanese variety Toyohomare (*I* allele) containing ∼800 bases of the DnaJ gene next to two tail-to-tail oriented CHS genes followed by CHS1 (Kasai et al., 2007; Senda et al., 2012). In the *i^k^* allele, recombination placed a partial P450 gene fragment upstream of the two inverted repeat CHS genes (Figure 4C, D). Expression data for the *i^i^* allele showed that while DnaJ is expressed similarly in hilum and seed coat tissues, P450 gene expression is significantly lower in the hilum (Figure 5). This pattern explains the dominance of the I allele, where DnaJ-driven CHS siRNAs are expressed in both tissues, over *i^i^* and *i^k^* alleles, which restrict CHS siRNA expression to the seed coat. The *i* allele is recessive to all other alleles because the allele does not express CHS siRNAs in either tissue.

The soybean whole-genome duplication generated inverted repeat-rich regions in sRNA loci (neo-functionalization), driving cis-regulatory mutations and tissue-specific expression of sRNA (regulatory subfunctionalization). Duplication and speciation of sRNA genes result in new targets, expression patterns, and functional divergence (Cui et al., 2017). The inverted duplication hypothesis suggests that plant sRNA genes originated from inverted duplications of their target genes (Allen et al., 2004; Fahlgren et al., 2007). In snapdragon, an inverted duplication led to the de novo production of functional sRNAs in snapdragon blooms, illustrating how structural genome changes generate new sRNA loci, facilitating neofunctionalization (Bradley et al., 2017). Similarly, in monkeyflowers (*Mimulus*), an inverted repeat at the YUP locus produces phased siRNAs that regulate floral carotenoid pigmentation, contributing to phenotypic diversification and speciation (Liang et al., 2023).

Tissue-specific gene expression, a fundamental aspect of cellular differentiation and organismal development, is shaped by mechanisms such as gene duplication, subfunctionalization, and regulatory element evolution (Lan and Pritchard, 2016; Sonawane et al., 2017). The evolution of tissue-specific expression patterns involves cis-regulatory elements that act locally in an allele-specific manner, enabling tissue- and temporal-specific expression with reduced pleiotropy, thereby modulating evolutionary change (Hill et al., 2021). In cichlid fish, about 20% of duplicate gene pairs acquired new tissue-specific functions driven by minimal regulatory mutations and accelerated sRNA evolution (Brawand et al., 2014). Similarly, the Arabidopsis miR166 family regulating meristem development, emerged through genome alterations followed by tissue-specific subfunctionalization (Maher et al., 2006).

Repeat-rich regions are powerful drivers of genomic structural changes that can influence important traits, yet they are often overlooked due to the challenges associated with investigating them. While seed coat color is a visually obvious trait influenced by repeat regions, it is likely that other traits are similarly controlled by repeat-rich regions. For example, soybean cyst nematode (SCN) resistance is primarily governed by the rhg1 locus, which contains multiple genes. SCN resistance has been shown to depend on the number of tandem repeats in this region (Cook et al., 2012). Although less prevalent in animals and humans, gene duplications and rearrangements can have significant deleterious effects, often contributing to diseases (Ting et al., 2004; Zhang et al., 2009). In soybeans, genome rearrangements produce nearly 100% identical repeat regions (Supplemental Figure), making genome assembly and structural or gene expression analyses more challenging (Schmutz et al., 2010; Wm82a1 and Wm82a2, soybase.org). The chimeric gene Glyma.08G110380, which includes partial sequences from subtilisin, CHS1, and CHS3, exemplifies these difficulties. Its 100% identical repeat regions (Supplemental Figure 4) can lead to over- or under-representation of aligned reads, complicating accurate quantification of gene expression. Given these limitations, aligning reads through the intergenic regions between the partial subtilisin fragment and the antisense CHS1 and CHS3 genes can provide a more reliable method for quantifying chimeric gene expression (Figure 5 and Supplemental Figure 5).

In summary, the whole genome duplications and subsequent structural rearrangements have shaped the soybean genome, resulting in domestication traits like seed coat pigmentation. Our study demonstrates how repeat-rich regions and cis-regulatory mutations drive tissue-specific gene expression and trait development. Advancements in sequencing technologies, particularly long-read methods, offer powerful tools to explore repetitive genomic regions, uncovering their broader contributions to evolutionary processes and crop improvement.

## METHODS

### Plant Materials

Soybean (*Glycine max*) lines used in this study are inbred and homozygous for the indicated loci. Lines are available through GRIN (Germplasm Resources Information Network). The full list of lines used in this report is shown in Supplemental Data Set 1. Plants were grown under standard greenhouse lighting and temperature conditions.

### DNA extractions for PCR

DNA was extracted from shoot tips of young plants that had been freeze-dried. Standard methods were used as in Cho et al., 2017, except that DNAs used for PCR screening only were not processed by phenol and chloroform extractions. Extraction buffer containing 100 mM Tris-HCl, pH 8.0, 500 mM NaCl, 50 mM EDTA, 1% SDS, 10 mM phenanthroline, 10 mM mercaptoethanol was added to tissue that had been ground and incubated at 68°C for 10 min. After addition of 5M potassium acetate, samples were incubated on ice for 15 min. Samples were centrifuged for 10 min and the supernatant transferred to cold isopropanol. The precipitated DNA was collected by centrifugation for 5 min in a microfuge and the supernatant decanted and redissolved in a small amount of TE (10 mM Tris, 1 mM EDTA, pH 7.5). After reprecipitation with 1/10 volume of 3 M sodium acetate and 2 volumes of isopropanol, the DNA was collected by centrifugation and dried with a Savant SpeedVac. The DNA pellet was redissolved in TER (10 mM Tris, 1 mM EDTA, pH 7.5, with 20 g/ml RNase) and incubated at 37°C for 30 min. DNA concentration was determined by Nanodrop (ThermoFisher) measurements.

### PCR Primer Selections and Amplifications

We chose primers that were unique in the genome Wm82.a2.v1 reference genome (according to Phytozome, Joint Genome Institute, Department of Energy), or had sufficient mismatches to similar genes on Chr05 (checked by eye with MultAlin, Corpet 1988), and caused no self-dimers or cross-dimers to each other (checked with ThermoFisher Multiple Primer Analyzer). Primers were designed to amplify almost the entirety of the hypothetical gene, Glyma.08G110600. The forward primer 71F (ACTCCGGCATGTCCACAGAAT) is found in the 5’UTR of the gene and the reverse primer 480DR (CTCCCATTATCATCACATTGG) is found approximately 500 nt downstream of the 3’UTR of the gene. The amplicon between 71F and 480DR (inclusive of primers) is 2764 nt. The primers have mismatches and insertions compared to Glyma.05G153200, a similar gene on Chr05 as shown in Supplemental Figure 1. For the RNA Recognition Motif (RRM) gene, Glyma.08G109600, primers designed to directly amplify the gene were not successful. Thus, primers were designed to amplify a region upstream of the 5’UTR of the gene. The forward primer Up2 (TTGATGCCCAGCAGTTGCTAG) and the reverse primer Up14R (AACCCAATTTAGGTATATCAG) are both found within 5000 nt upstream of the 5’UTR. The amplicon between the primers (inclusive) is 2909 nt. The primers have mismatches and insertions compared to the upstream region of Glyma.05G152900, a similar RRM gene on Chr05 as shown in Supplemental Figure 1. The presence or absence of this amplicon was inferred to indicate the presence or absence of the RRM gene in PCR reactions. To amplify the 7.9 kb fragment characteristic of the NAHR deletion lines as in W55, we used the same primers described in Cho et al, 2019. Primers BC1F (CTATAATCATATCTATGGACTCTCCCTCAC) and BC2R (AACTTTAACCTCTCATGTAAGGAAACAAATAAC) are found in the 5’-most epimerase gene Glyma.08G109100 and in the 5’ region of the galactosidase gene Glyma.08G111000, respectively. These primers span a deletion of approximately 138 kb and only amplify the 7.9 kb fragment when this region has been deleted, as it has in certain Williams *i* mutants such as W55. Due to the deletion these varieties are normally missing the hypothetical and RRM genes that would be amplified by the other primers mentioned above. Polymerase chain reactions were performed in 50 *μ*l volume reactions in 0.2 ml Corning Axygen thin wall PCR tubes (#14-222-262, Fisher Scientific). For the amplifications of the hypothetical and RRM genes, 1 *μ*l of 0.5 *μ*g/*μ*l DNA was used and 1 *μ*l of each primer at 10 *μ*M per reaction. To this was added 5 *μ*l 10x Ex Taq Buffer (contains 20 mM Mg2+), 4 *μ*l dNTP mix (2.5 mM each), and 0.25 *μ*l Takara Ex Taq polymerase (all part of the Takara Ex Taq kit, RR001A), per reaction. Amplifications were performed in a PTC-200 DNA Engine from MG Research using the following cycle times: initial denaturation at 94°C for 1 min, followed by 30 cycles total of denaturation at 94°C for 30 sec, annealing at 55°C for 1 min, and elongation at 72°C for 1 min; ending with an extension at 72°C for 2 min. For the amplification of the 7.9 kb fragment, reagents were the same except 2 *μ*l of 0.5 *μ*g/ul DNA was used per reaction. A slightly different amplification program was used, with an initial denaturation at 94°C for 1min, followed by 30 cycles total of denaturation at 98°C for 10 sec, annealing at 55°C for 1 min, and elongation at 68°C for 9 min; ending with an extension at 72°C for 10 min.

### Illumina Short Read Sequencing and Data Analyses

Sequencing of the W44 mutation was conducted by the DNA Service facilities of the University of Illinois with sheared DNAs of average size 500 bp. Libraries were prepared with Kapa Library Construction Kits and quantitated by qPCR and sequenced using a TruSeq SBS sequencing kit version 1 in one lane of an Illumina HiSeq2500 instrument from each end yielding 322 million total reads of 100 nt. Fastq files were generated with the software Casava 1.8.2 (Illumina). Alignments of short reads were made as previously indicated (Cho et al., 2019) using Bowtie v2 (Langmead et al., 2012) to the Williams reference genome assembly and then displayed using the Integrative Genomics Viewer (Robinson et al., 2011).

### High Molecular Weight DNA Preparation and Sequencing and Assembly of Long Reads

For long read sequencing with PacBio, soybean plants were grown for 17 to 19 days in 5-inch pots under standard greenhouse conditions with 14 hrs of natural or supplemental light with day temperatures controlled between 25-28°C and night between 19 to 21°C. The pots were then transferred to the dark for a 48-hr treatment to minimize chloroplast DNA accumulation prior to harvesting the shoot tips and young biofoliate and trifoliate leaves. The leaves were frozen in approximately 5 to 10-gram aliquots in heat sealed Kapak pouches in liquid nitrogen and stored at −80 °C. Nuclei were isolated in a large-scale prep from up to 40 grams of tissue, and high molecular weight DNA was extracted from one sample (UC8) by Polar Genomics, Ithaca NY, yielding 550 *μ*g of high molecular weight (HMW) DNA between 100-150 kb as judged by pulsed field electrophoresis and concentration measurements with a Qubit instrument. Since only 5 *μ*g of HMW DNA is needed for sequencing, a smaller scale procedure was developed by the DNA Sequencing Services of the Carver Biotechnology Center at the University of Illinois using 2 grams of leaves, which typically yielded between 10 and 15 *μ*g of HMW DNA between 100 and 200 kb for each of the four other lines (Richland, T157, UC3, and W44). The procedure consisted of nuclei isolation following the LN2 Plant Tissue Protocol Document ID: NUC-LNP-001 from PacBio/Circulomics with the exception of maintaining the samples on ice at all times and filtering the lysate with 100 *μ*m and 44 *μ*m size mesh. DNA isolation was performed with the MagMax Plant DNA kit (ThermoFisher, catalog number A32549) kit.

Library preparation and long read sequencing were also conducted by the University of Illinois Sequencing Center. The HMW DNA was sheared with a Megaruptor 3 (Diagenode.com) to an average fragment length of 13 kb. Sheared DNAs were run on the BluePippin (Sage Science) using a 0.75% agarose cartridge for size selection of fragments > 10kb in length. Size selected DNAs were converted to a library with the SMRTBell Express Template Prep kit 2.0. Each library was sequenced on 2 SMRTcells on a PacBio Sequel IIe using the CCS sequencing mode and a 30h movie time. CCS analysis was done using SMRTLink V10.1 using the following parameters: ccs --min-passes 3 --min-rq 0.99. Resulting raw reads of from 40 to 77 Gb for each variety were assembled by the computing facilities of the Carver Biotechnology Center for High Performance Computing in Biology using the software Hifiasm version 0.16.1. Metrics from the SMRT cell yield and assembly for each variety both before and after removal of mitochondrial or other contaminant contigs by the Foreign Contamination Screen tool suite (Astashyn et al., 2023) used by the National Center for Biotechnology Information are shown in Supplemental Data Set 2. Assignment of the genes in the *I* locus regions of each assembly were conducted with local BLAST alignments as described in Supplemental Data Sets 3-9. Could put some of the discussion of these here rather than in the Results.

### RNA-Seq and Small RNA-Seq and Data Analyses

Total RNA was isolated by standard methods as previously described (Wang and Vodkin, 1994) using phenol and chloroform extractions followed by lithium precipitations. For small RNA isolations, the lithium chloride precipitations were omitted. Preparation of libraries and Illumina sequencing were conducted by the University of Illinois DNA Services Center as described generally in (Cho et al, 2017 and 2019) and with specific kits and Illumina machines as described in the more detail in the Gene Expression Omnibus (GEO) accession entries for each library.

After adapter trimming, the majority of small RNAs were generally in the range of 18 to 25 nt and were analyzed using Bowtie v.1 (Langmead et al., 2009) for alignments against all 86,256 Glyma model transcripts including splice variants from the Wm82.a4 reference genome. Positive hits, negative hits, and total hits were aggregated per model and normalized per million total reads for comparison between the same gene models. RNA-seq datasets were similarly analyzed with Bowtie v.1 with up to 3 mismatches allowed against all transcripts of the reference genome Wm82a.2 and hits per transcript were normalized to RPKM values (reads per million per kilobase of model) as described (Mortazavi et al., 2008).

Alternatively, the RNA-seq data were aligned with Bowtie v.1 to the 186,027 bp genomic region from the reference genome Wm82a.4 Gm08:8360676-8546687 surrounding the *i^i^* allele with Bowtie v1 (Langmead et al., 2009) with no mismatches allowed. We used the commands (-best –strata -M 1) that chose one among multiple valid alignments, which had a 100% match, and randomly distributed it (Treangen and Salzberg 2011). Output files in SAM format were converted to BAM format and sorted with the Samtools program (Li et al., 2009). The Integrative Genomics Viewer (IGV) (Robinson et al., 2011) was used to convert from BAM to TDF format and display normalized data (count at base x one million/total number of reads).

### Accession Numbers

The genome assembly contigs and PacBio raw data can be accessed at the National Center for Biotechnology Information with Bioproject accessions and Whole Genome Shotgun accessions numbers as follows: for UC3, PRJNA987730 and JAVIAI000000000; for W44, PRJNA988802 and JAVIAJ000000000; for Richland, PRJNA989342 and JAVIAK000000000; for T157, PRJNA990340 and JAVIAL000000000; and for UC8, PRJNA990476 and JAVIAM000000000. Illumina short read resequencing data for the W44 mutation is under PRJNA505340. The 38 new smRNA-seq samples and 50 new RNA-seq entries for various tissues or stages of development can be found in Gene Expression Omnibus with series accessions GSE239923, GSE239484, and GSE41201. Supplemental Data Sets 10-14 have the complete information including accession numbers for all smRNA-seq or RNA-seq samples used in this report.

## SUPPLEMENTAL DATA

**Supplemental Data Set 1.** Complete Information for Soybean Mutant Lines from the USDA Soybean Germplasm Screened by PCR or other Whole Genome Sequencing Technologies.

**Supplemental Data Set 2.** Metrics from Sequencing and Assembly of Long Reads.

**Supplemental Data Set 3**. Placement of *CHS* Genes in the *I* locus Regions of the Assembled Genomes.

**Supplemental Data Set 4.** Annotation of the SHMT to Gal Regions of the Five PacBio Sequences.

**Supplemental Data Set 5.** Raw Read Retrieval from UC3, *i* Mutation with Black Seed Coats.

**Supplemental Data Set 6.** Raw Read Retrieval from W44 line, Williams i Mutation with Black Seed Coats.

**Supplemental Data Set 7.** Raw Read Retrieval from Richland, *I* with Yellow Seed Coats.

**Supplemental Data Set 8.** Raw Read retrieval from T157, *i* Mutation with Imperfect Black Seed Coats.

**Supplemental Data Set 9.** Raw Read Retrieval from the UC8 Isoline Homozygous for the Black Saddle *i^k^* Allele.

**Supplemental Data Set 10.** RNA-seq Data Contrasting Hila versus Seed Coat Without the Hila from Williams *i^i^* genotype.

## ACKNOWLEDGEMENTS

We are grateful to Dr. Alvaro Hernandez and staff of the University of Illinois High-Throughput Sequencing Unit of the Biotechnology Center for the PacBio and Illumina genome, mRNA, and small RNA sequencing. We are grateful for the services provided by Kimberly Walden with the University of Illinois High Performance Computing in Biology (HPCBio) with the Biotechnology Center for application of the Hifiasm/0.16.1 genome assembly algorithm for the long-read sequencing data. The research was supported by the University of Illinois Foundation and United Soybean Board.

## AUTHOR CONTRIBUTIONS

YBC, SIJ, and LOV collected tissue samples, performed experiments, analyzed, and interpreted data, and drafted results and manuscript sections. SIJ conducted PCR analyses, YBC performed genome and RNA-seq alignments, LOV performed local BLAST genome and RNA analyses. LOV designed approach, led, and coordinated the project. All authors have read and approved the manuscript.

## DISCLOSURE: USE OF LLM AI (ChatGPT)

This manuscript utilized ChatGPT solely for proofreading purposes. All citations were thoroughly verified by the authors.

