## Supplemental Figures and Tables for "Cis-regulatory Mutations Drive Tissue-specific Subfunctionalization of sRNA loci Regulating Soybean Seed Color During Domestication"

**Supplemental Figure 1.** PCR Results Distinguish Large NAHR Events from Smaller changes in Multiple Independent Self-Color Mutations from the Soybean Germplasm Collection.

**(A)** Diagram illustrating the structure of the *i^i^* allele of the *I* locus and non-allelic homologous recombination (NAHR) events known to delete 138 kb between the most distant *CHS* gene (adapted from Cho et al., 2019). The gene models from the reference genome Williams 82 (*i^i^*) Wm82a.4 are shown. Williams 82 is a disease resistant inbred isoline of Williams (W43). Numbers are kb (kilobases) on chromosome 8. The red and yellow horizontal arrows represent regions of identical or nearly identical repeated sequences. Yellow curly brackets show the position of the *CHS* repeats of the *I* locus. Green and purple arrows show the positions of primer sets for epimerase (Glyma.08G109100) and a 5’ region of the galactosidase gene (Glyma.08G111000), respectively, and illustrate an enlargement of the 7.9 kb amplicon that results. The epimerase primer will match the repeat epimerase region but in the opposite direction as denoted by the dotted green arrow. Dotted lines illustrate that the recombination event occurred between the distal *CHS5* and *CHS1* genes to form a *CHS5:1* hybrid as revealed by sequencing the 7.9-kb amplicon in several black seeded mutations.

**(B)** Three representative gels of PCR reactions with different primer sets for the RRM gene (blue), hypothetical gene within the *I* locus region (orange), and the 7.9 kb amplicon discussed above that were used in screening 30 recessive *i* mutations that will distinguish the large NAHR events from other smaller changes. Seed pictures are representative of the color type. M are standard markers used in sizing. The primer sets are shown in Supplemental File 1 and Materials and Methods. The controls used in all gels are the previously characterized W55 and W130 which are independent mutations from the parent cultivar Williams (W43) as shown by the arrows as well as UC9 which is a mutation from the variety Clark 63 (UC7). Both had been characterized by amplicon sequencing of the 7.9-kb PCR fragment (Cho et al., 2019).

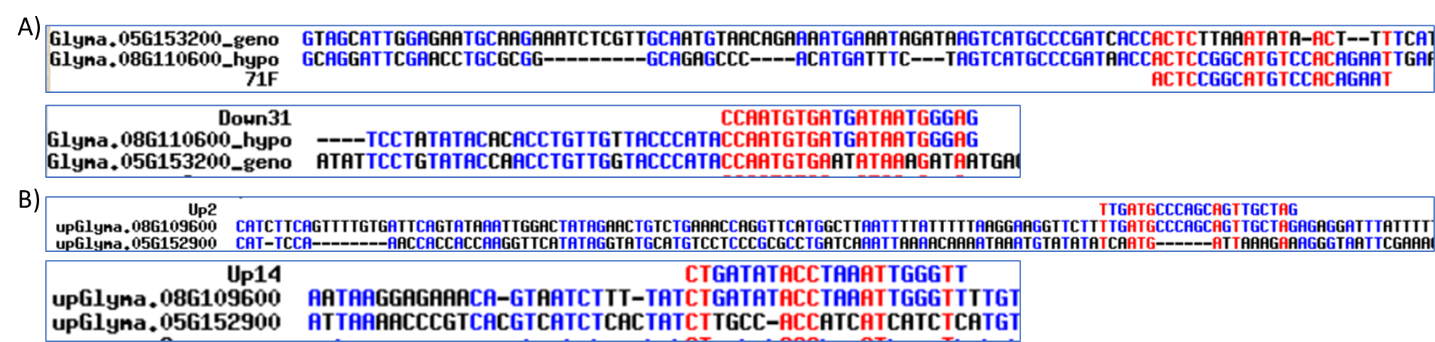

**Supplemental Figure 2.** Hypothetical and RRM Gene Primers Shown in Multalin v2 alignment.

Primers for the hypothetical and RRM genes compared to the intended gene regions on Chr08 and similar regions on Chr05. The primers match 100% to the intended gene regions on Chr08 but have mismatches and insertions compared to similar gene regions on Chr05. Sequences compared in Multalin. Red letters indicate a match across all three sequences, blue letters a match across two sequences, and black letters are unique to one sequence. Dashed lines indicate where bases have been inserted compared to the other sequence.

**(A)** Primers for the hypothetical gene Glyma.08G110600 compared to similar gene Glyma.05G153200. The primers are 71F (5’ACTCCGGCATGTCCACAGAAT3’) and 480DR (5’CCAATGTGATGATAATGGGAG3’, shown as reverse complement Down31). 71F matches to the 5’UTR of the gene and 480DR is the reverse complement matching downstream of the 3’UTR. The amplicon between 71F and 480DR (inclusive) is 2764 nt.

**(B)** Primers for the RNA Recognition Motif (RRM) upstream region of gene Glyma.08G109600 compared to similar gene Glyma.05G152900. The primers are Up2 (5’TTGATGCCCAGCAGTTGCTAG3’) and Up14R (5’CTGATATACCTAAATTGGGTT3’), shown as reverse complement Up14. Both match within 5000nt upstream (shown here for both genes) of the 5’UTR. They produce an amplicon of 2909 bp.

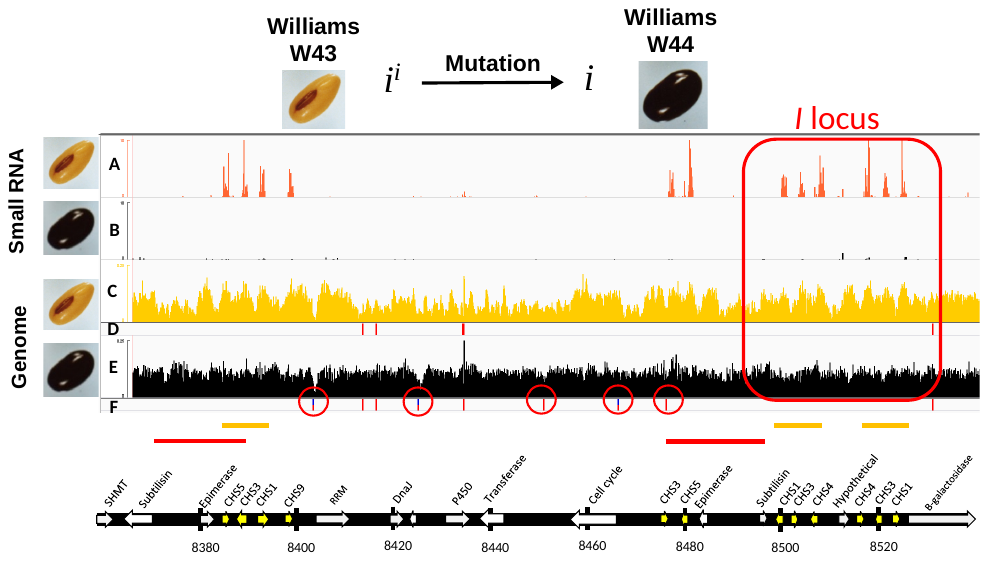

**Supplemental Figure 3.** Williams, W44, *i* Mutation, Abolishes Small RNA Production but Does Not Reveal any Structural Variation by Short-Read Alignments. Numbers are kb on chromosome 8.

**(A)** and **(B)** Alignments of small RNA-seq data [scale: 0-10] from 50-100 mg seed coats from Williams (W43) and Williams (W44).

**(C)** and **(D)** Alignments of short-read, whole genome sequencing data [scale: 0-0.25] from the Williams (W43) parent and SNP calls from the data (vertical red lines).

**(E)** and **(F)** Alignments of short-read, whole genome sequencing data [scale: 0-0.25] in the Williams (W44) black mutation and SNP calls from the data (vertical red lines).

Alignments were made to the reference Williams 82 genome with annotation shown below for the SHMT to galactosidase region. Normalized alignment coverage data [count at base × one million / total number of reads] are visualized by the Integrative Genomics Viewer (IGV) tool. The five circles indicate additional SNPs only in the Williams 44 mutant. The horizontal red lines or yellow lines indicate areas of exact or very similar repeated sequences. The region of the *I* locus is indicated by a red box.

**Supplemental Table 1**. Assembled Contigs Containing Chr 8 and Positions Around the I Locus

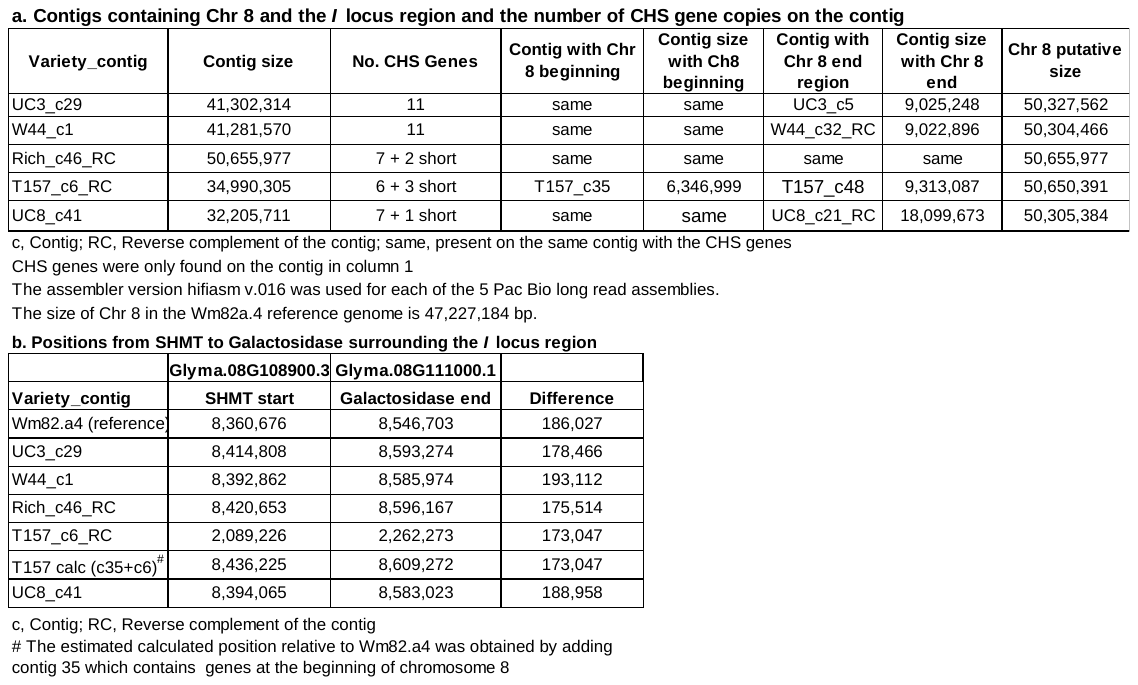

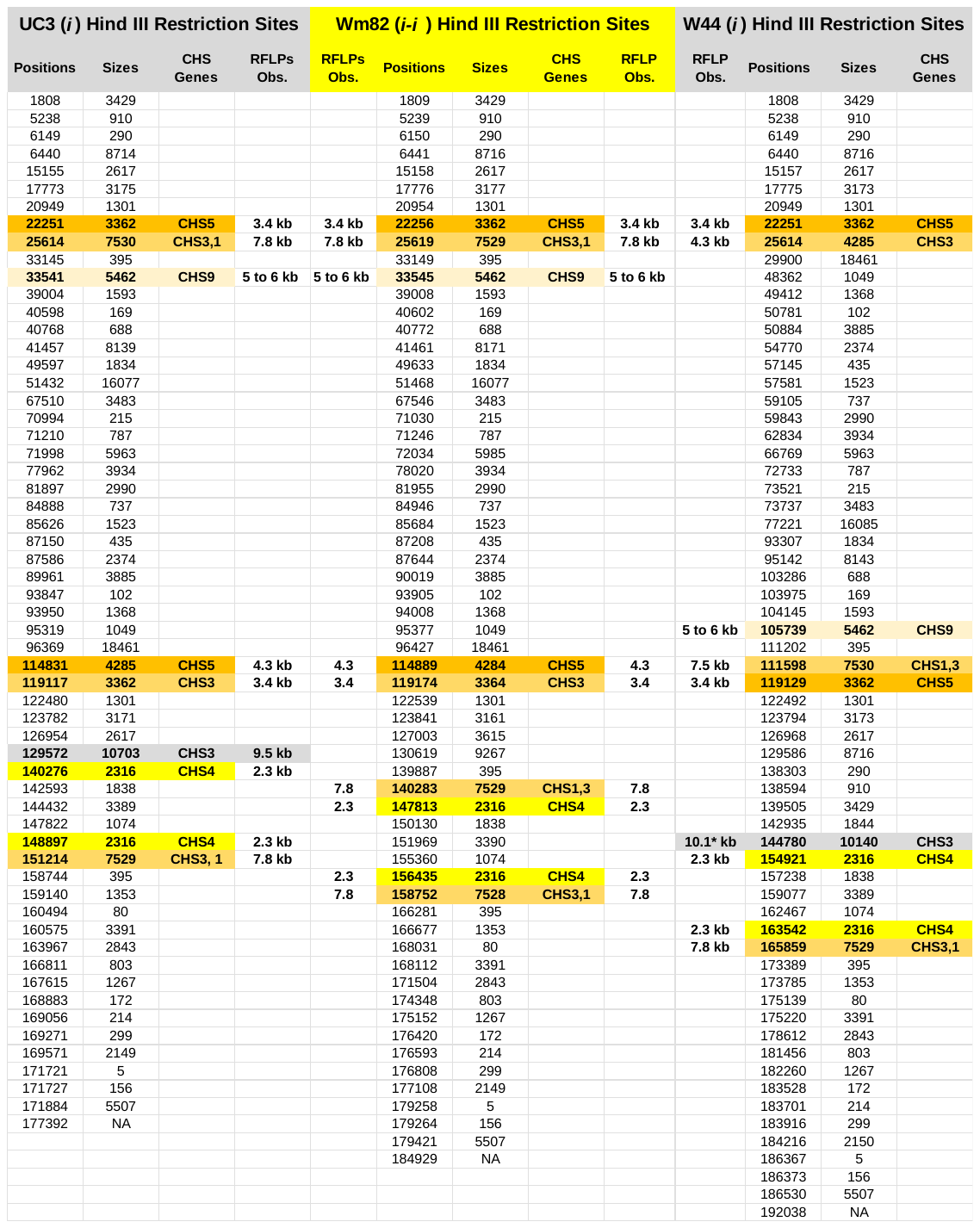
**Supplemental Table 2**. Variation in HindIII Restriction Sites in UC3 and W44 Mutant Lines. Sizes are compared to the Wm82a.4 genome. The region between SHMT to Gal was analysed. RFLPs are the observed sizes estimated in blots shown in Todd and Vodkin, 1996, that uses a coding sequence probe that would hybridize to all the CHS genes. An extra band estimated at 9.5 kb was observed in UC3 line. *indicates that a band at 10 kb was not observed in the in W44 line, likely from incomplete separation in the sample at the larger sizes. Incomplete digestion of the 7.5 and 2.3 kb bands can lead to shadows at sizes in the 9 kb to 10 kb range.

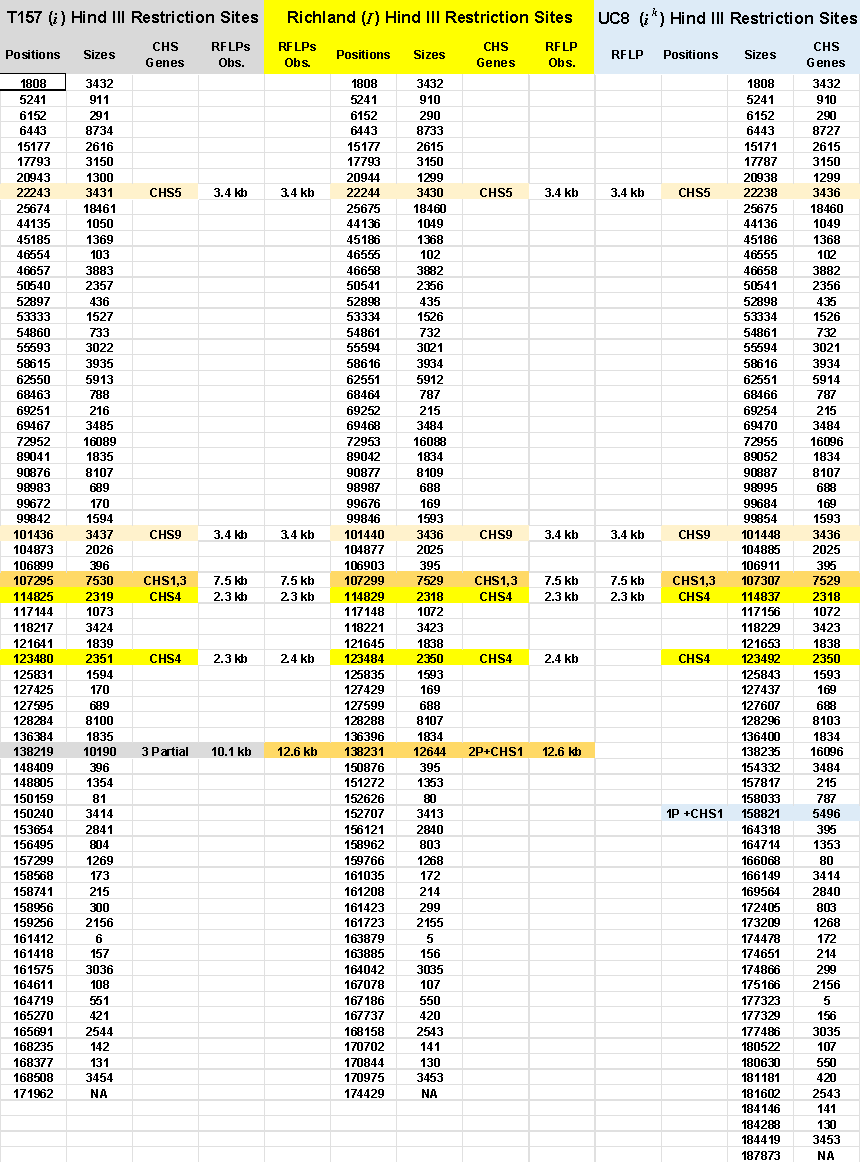

**Supplemental Table 3**. Variation in HindIII Restriction Sites in Richland, T57, and UC8. The region between SHMT to GAL was analysed. Obs, observed RFLPs are the estimated sizes in blots shown in Todd and Vodkin, 1996, for Richland and T157, that uses a coding sequence probe that would hybridize to all the CHS genes. UC8 had not been subjected to DNA blotting and sizes listed are the same as the sequence digest predicts.

**Supplemental Figure 4.** Comparison of Upstream Regions of the Partial Subtilisin *CHS* siRNAs and the Cognate Subtilisin Gene of Origin.

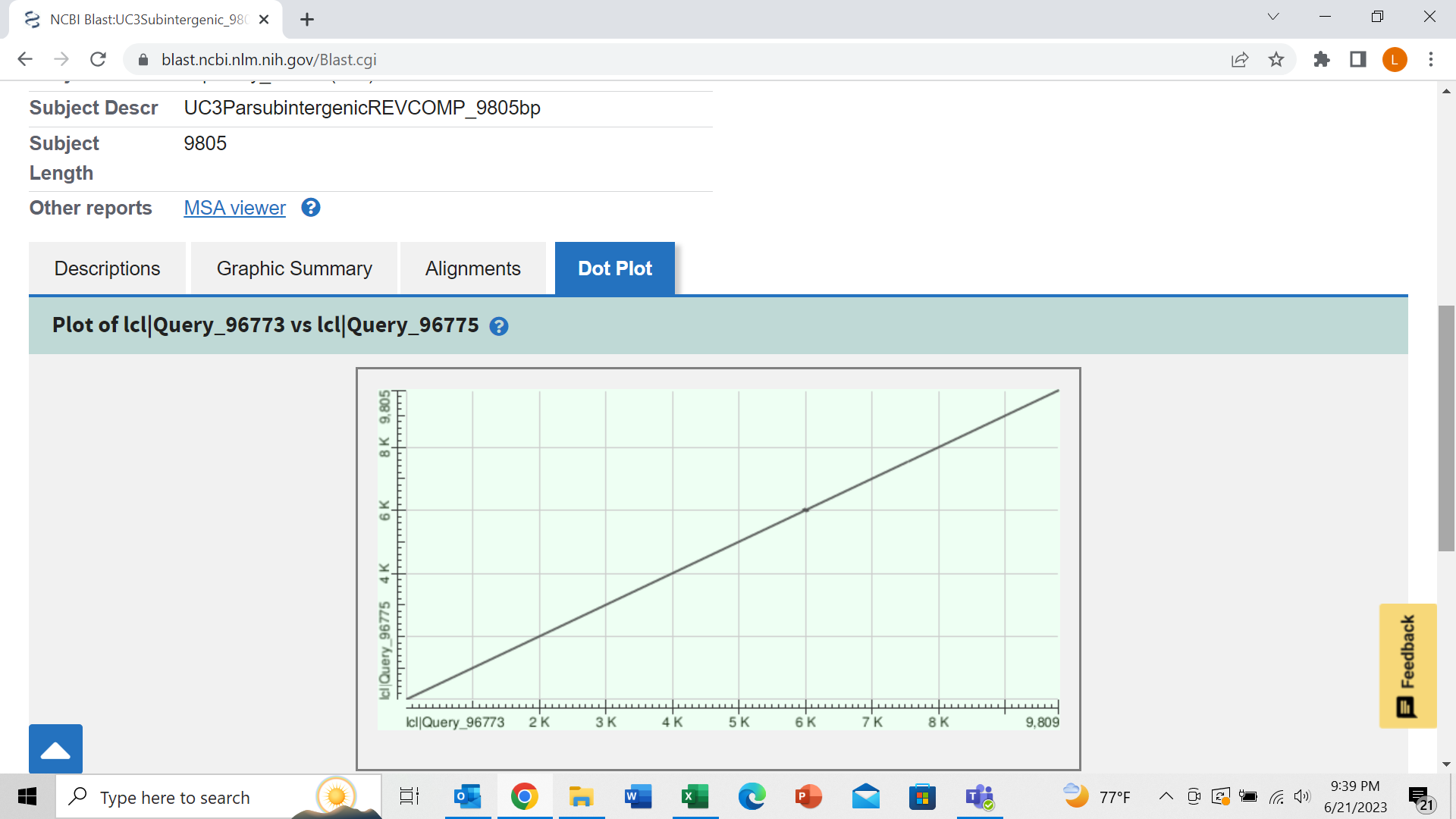

**A. UC3 (*i^i^* to *i* mutation) blast alignments.**

Dot plot of blast alignments of the 9805 bp intergenic region upstream of the partial subtilisin gene and the compared to the 9809 bp upstream region of its cognate origin gene (equivalent to Glyma.08G109000.1 in Wm82a.4). They are identical except for four ATAT bases within a far-upstream AT-rich region.

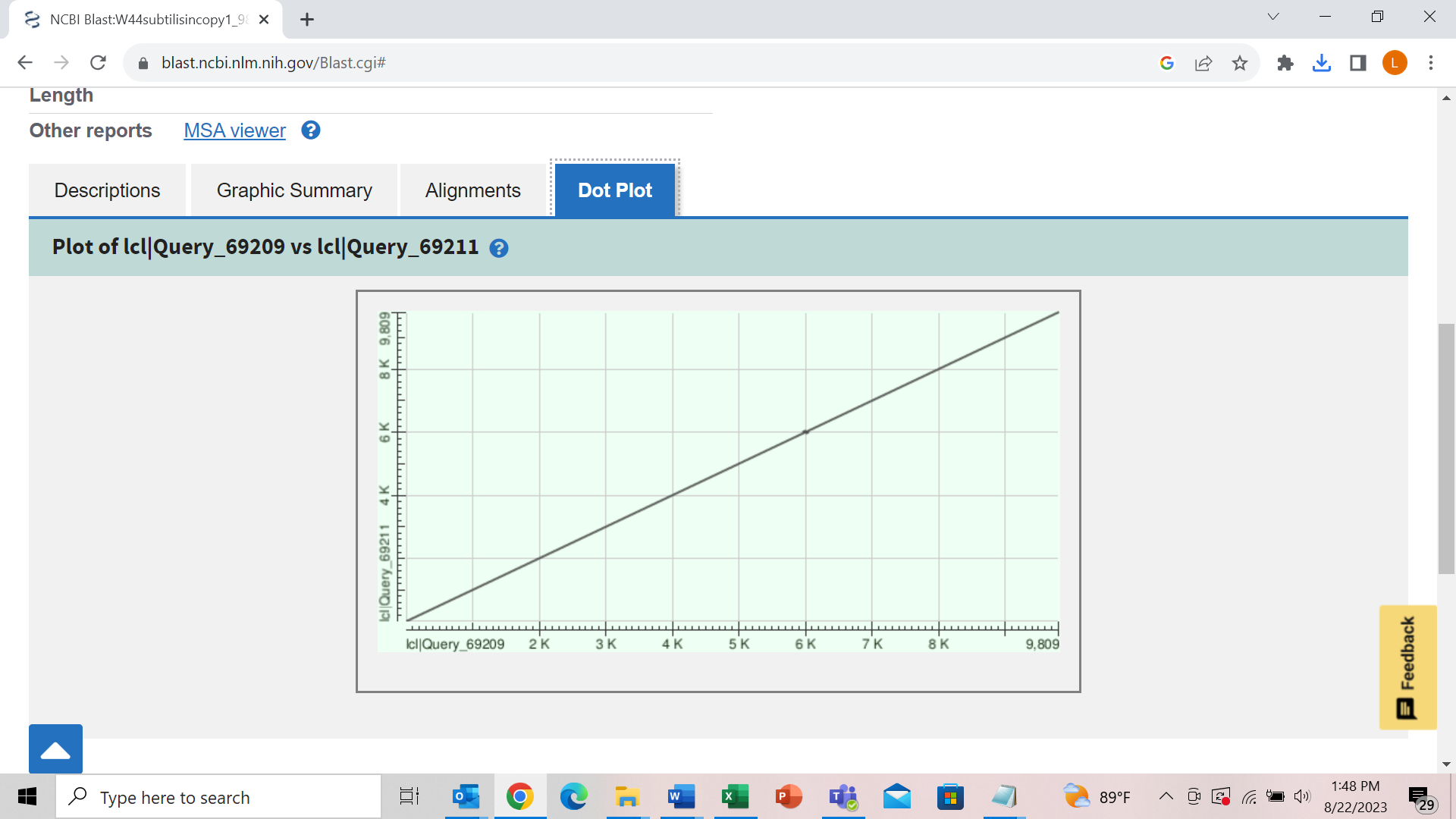

B. **W44 (*i^i^* to *i* mutation) blast alignments**.

Comparison of the two complete subtilisin 9808 bp gene copies of the W44 genome (*i^i^* to *i* mutation) which are identical.

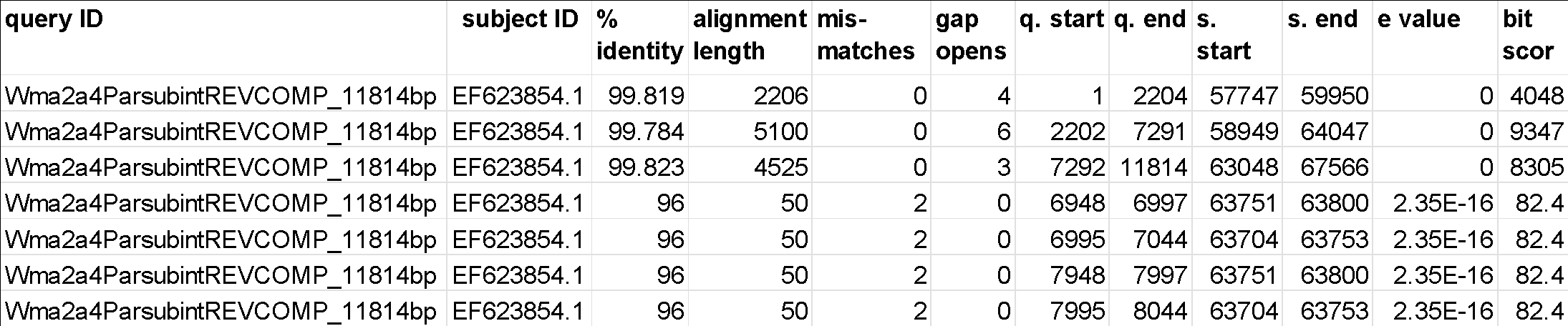

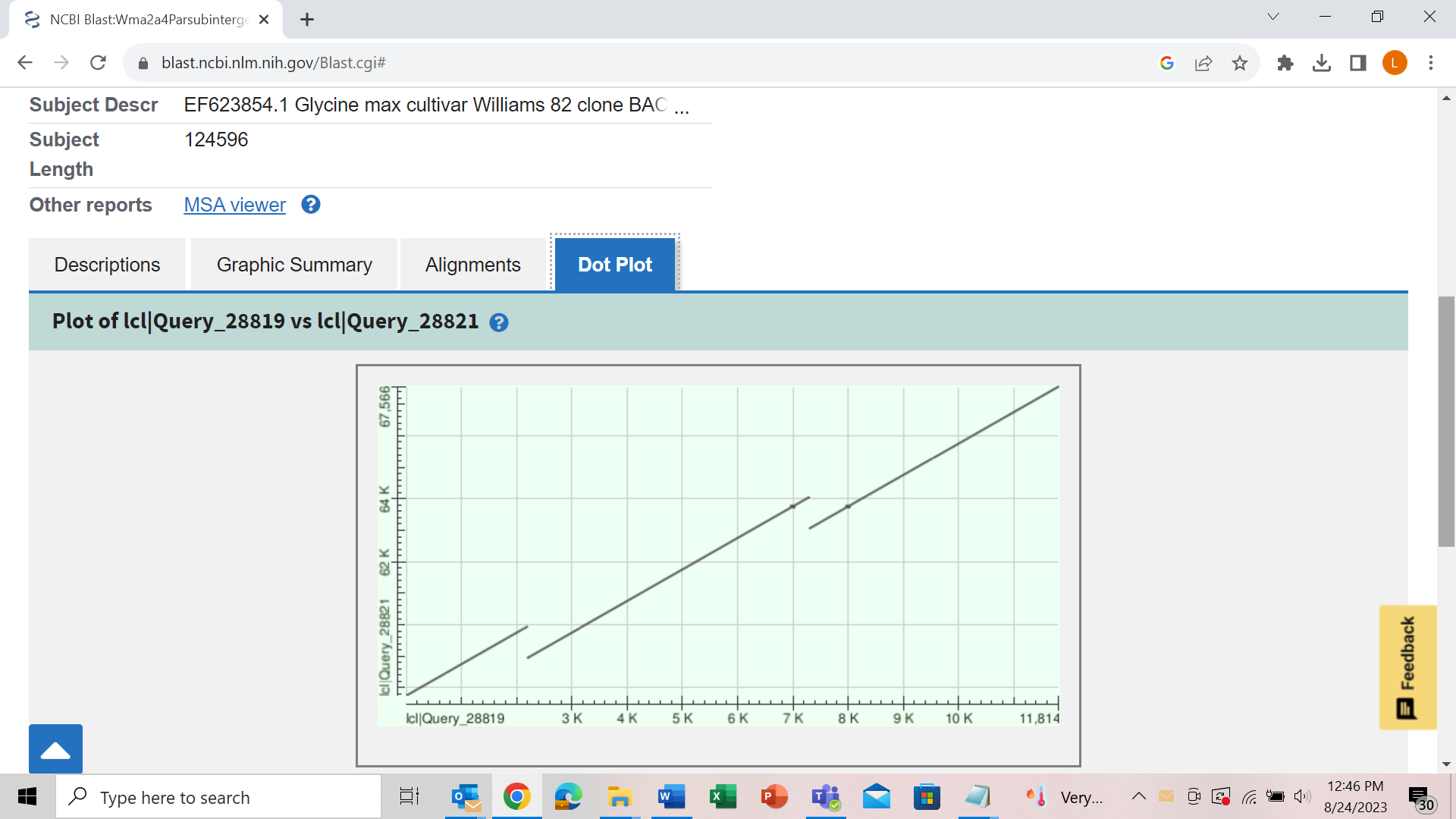

**C. The Wm82a.4 reference genome compared to Wm82 BAC sequences.**

The 11,814 bp region upstream of the partial subtilisin of the Wm82a.4 reference genome as query to the same region of BAC77G7-a (accession number EF623854.1) also isolated from Williams 82 with the *i^i^* allele. The alignment hit table and the dot plot show that the Williams 82 BAC77G7-a does not have the two internally repeated areas that the Wm82a.4 reference genome assembly has. Thus, these repeated regions are likely false automated assembly errors in this region of the Wm82a.4 reference genome.

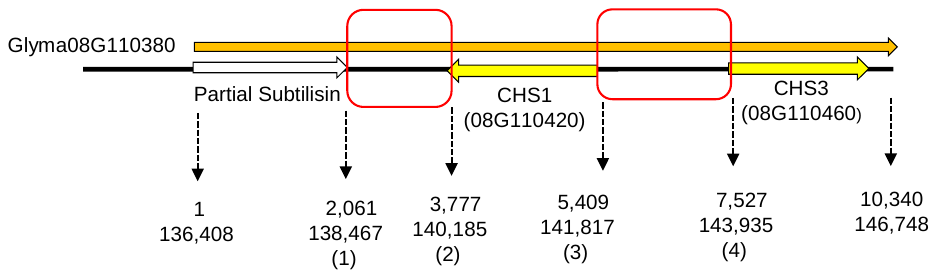

**Supplemental Figure 5.** Chimeric RNA-seq Alignments in the Subtilisin-Driven *i^i^* Allele.

**(A)** Diagram of the structure of the Glyma08G11038 (10,340 bp) gene call in the Wm82a.4 genome which is the origin of the double-stranded RNA transcripts in *i^i^* genotypes. It is an automated gene call and starts with the partial subilisin gene from position 1 to 2061 which is four of the 11 exons found in the cognate subtilisin gene. It then places the two authentic *CHS1* and *CHS3* genes within a large intron and terminates with 378 bases of similarity to the intergenic space between *CHS3* and *CHS4*. The relative positions of the junction sequences (red box areas) between the partial subtilisin, antisense *CHS1*, and sense *CHS3* genes are shown for the Glyma model and for the positions on the SHMT to Gal fasta file used for RNA-seq alignments. As shown in Figures 6A and 8B, most reads will come from the cognate subtilisin or repetitive CHS genes, but some low level of transcript reads appear to hit intergenic regions that cross junction sequences a, b, c, and d.

**(B)** Junction sequences in RNA-seq Bowtie alignments shown in Figure 6A for sample R52 (Williams 25-50 mg yellow seed coats of *i^i^* genotype).

**Junction 1:** The bases highlighted in blue are the end of the partial subtilisin hit region at nucleotide position 2061 of the complete subtilisin gene at position 138,467 in the SHMT to Gal fasta sequence of the reference Wm82a.4 genome sequence. These reads are chimeras between the partial subtilisin and the intergenic region separating it from the antisense *CHS1* gene. All reads are 75 bp.

GAGGGAGAAGGGGTAGAAGTAGAATCTCCATTTTACAATATACTTATCTTATTGACATCGGATCCTATACTTAAA

AGGGAGAAGGGGTAGAAGTAGAATCTCCATTTTACAATATACTTATCTTATTGACATCGGATCCTATACTTAAAT

AGAAGGGGTAGAAGTAGAATCTCCATTTTACAATATACTTATCTTATTGACATCGGATCCTATACTTAAATATAA

GGTAGAAGTAGAATCTCCATTTTACAATATACTTATCTTATTGACATCGGATCCTATACTTAAATATAAATATCT

AGTAGAATCTCCATTTTACAATATACTTATCTTATTGACATCGGATCCTATACTTAAATATAAATATCTAGTAGA

GTAGAATCTCCATTTTACAATATACTTATCTTATTGACATCGGATCCTATACTTAAATATAAATATCTAGTAGAA

GTAGAATCTCCATTTTACAATATACTTATCTTATTGACATCGGATCCTATACTTAAATATAAATATCTAGTAGAA

AGAATCTCCATTTTACAATATACTTATCTTATTGACATCGGATCCTATACTTAAATATAAATATCTAGTAGAAGT

AGAATCTCCATTTTACAATATACTTATCTTATTGACATCGGATCCTATACTTAAATATAAATATCTAGTAGAAGT

AGAATCTCCATTTTACAATATACTTATCTTATTGACATCGGATCCTATACTTAAATATAAATATCTAGTAGAAGT

TCCATTTTACAATATACTTATCTTATTGACATCGGATCCTATACTTAAATATAAATATCTAGTAGAAGTTCTTAA

TCCATTTTACAATATACTTATCTTATTGACATCGGATCCTATACTTAAATATAAATATCTAGTAGAAGTTCTTAA

TTACAATATACTTATCTTATTGACATCGGATCCTATACTTAAATATAAATATCTAGTAGAAGTTCTTAAGTTCCT

The IGV display below shows some of the reads (gray boxes) from the alignments for R52 that align uniquely because they are chimeras and do not match other regions of the genome as the upstream complete subtilisin gene or other CHS genes. The Bowtie alignments were conducted with no mismatches allowed and the best strata command lines to differentiate uniquely mapped reads versus those that match more than one location.

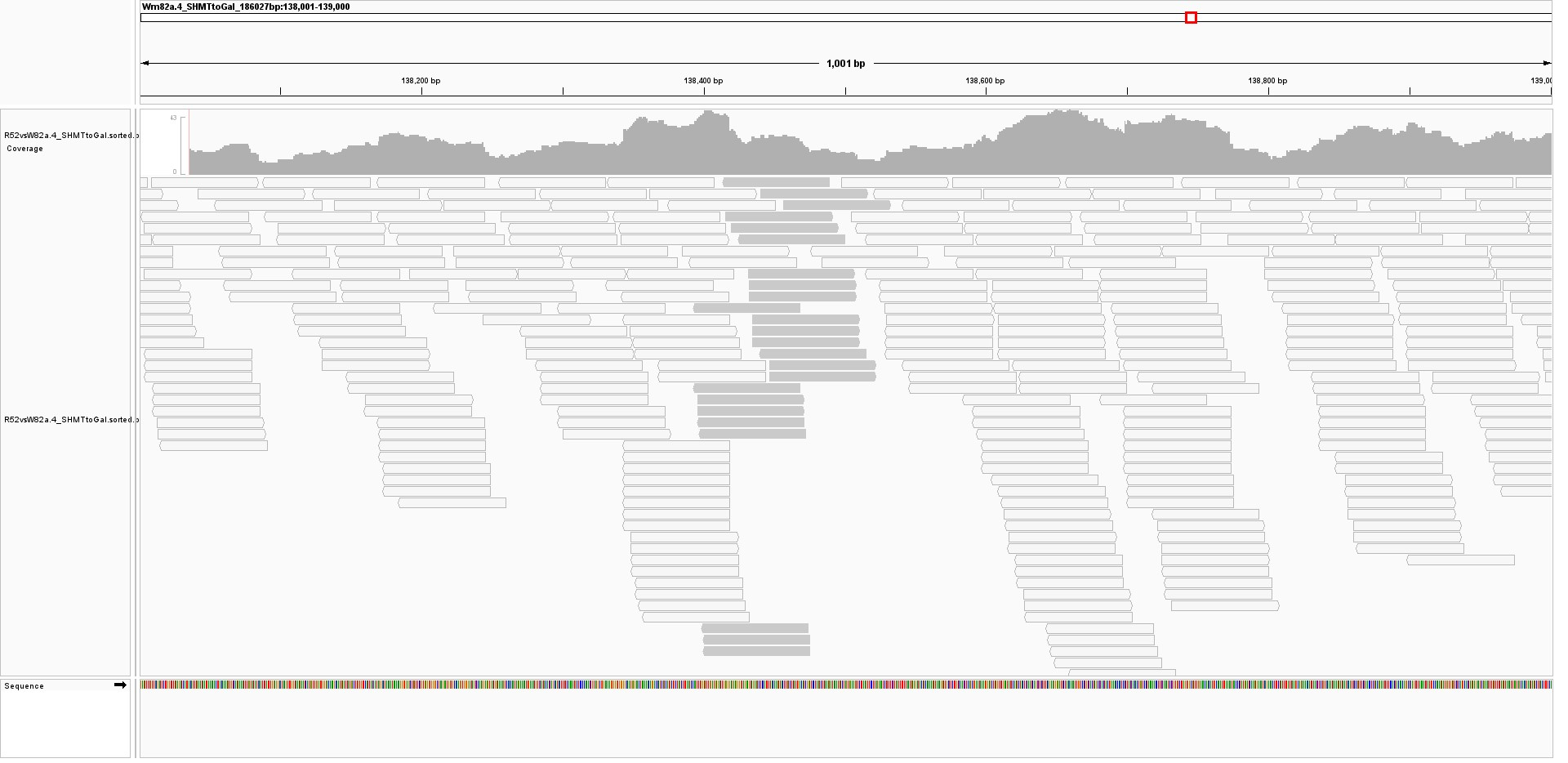

**Junction 2:** The bases highlighted in green are the first bases of the antisense *CHS1* gene Glyma.08G110420.1 gene showing the some of the chimeric RNA-seq reads connecting the preceding intergenic region to the *CHS1* gene.

CATACACAACACTCTTTACACAATTAGCCCATAAGTAAACCTGGTTCGAGACAGTTCTATAGTTCAATTTATCAT

TTTACACAATTAGCCCATAAGTAAACCTGGTTCGAGACAGTTCTATAGTTCAATTTATCATCAATCATAGAGAAT

TTTACACAATTAGCCCATAAGTAAACCTGGTTCGAGACAGTTCTATAGTTCAATTTATCATCAATCATAGAGAAT

TTTACACAATTAGCCCATAAGTAAACCTGGTTCGAGACAGTTCTATAGTTCAATTTATCATCAATCATAGAGAAT

ACACAATTAGCCCATAAGTAAACCTGGTTCGAGACAGTTCTATAGTTCAATTTATCATCAATCATAGAGAATCTT

CACAATTAGCCCATAAGTAAACCTGGTTCGAGACAGTTCTATAGTTCAATTTATCATCAATCATAGAGAATCTTT

AATTAGCCCATAAGTAAACCTGGTTCGAGACAGTTCTATAGTTCAATTTATCATCAATCATAGAGAATCTTTCAA

TTAGCCCATAAGTAAACCTGGTTCGAGACAGTTCTATAGTTCAATTTATCATCAATCATAGAGAATCTTTCAATG

CATAAGTAAACCTGGTTCGAGACAGTTCTATAGTTCAATTTATCATCAATCATAGAGAATCTTTCAATGAAGATA

ATAAGTAAACCTGGTTCGAGACAGTTCTATAGTTCAATTTATCATCAATCATAGAGAATCTTTCAATGAAGATAA

CCTGGTTCGAGACAGTTCTATAGTTCAATTTATCATCAATCATAGAGAATCTTTCAATGAAGATAAAATTGGACA

CCTGGTTCGAGACAGTTCTATAGTTCAATTTATCATCAATCATAGAGAATCTTTCAATGAAGATAAAATTGGACA

CCTGGTTCGAGACAGTTCTATAGTTCAATTTATCATCAATCATAGAGAATCTTTCAATGAAGATAAAATTGGACA

GTTCGAGACAGTTCTATAGTTCAATTTATCATCAATCATAGAGAATCTTTCAATGAAGATAAAATTGGACAAAAG

GTTCGAGACAGTTCTATAGTTCAATTTATCATCAATCATAGAGAATCTTTCAATGAAGATAAAATTGGACAAAAG

GTTCGAGACAGTTCTATAGTTCAATTTATCATCAATCATAGAGAATCTTTCAATGAAGATAAAATTGGACAAAAG

GTTCGAGACAGTTCTATAGTTCAATTTATCATCAATCATAGAGAATCTTTCAATGAAGATAAAATTGGACAAAAG

GTTCGAGACAGTTCTATAGTTCAATTTATCATCAATCATAGAGAATCTTTCAATGAAGATAAAATTGGACAAAAG

TCGAGACAGTTCTATAGTTCAATTTATCATCAATCATAGAGAATCTTTCAATGAAGATAAAATTGGACAAAAGAT

CGAGACAGTTCTATAGTTCAATTTATCATCAATCATAGAGAATCTTTCAATGAAGATAAAATTGGACAAAAGATA

**(C)** Junction sequences viewed by the IGV browser display for RNA-seq from different tissues. The light gray bars represent chimeric sequences between between the partial subtilisin and the intergenic region.

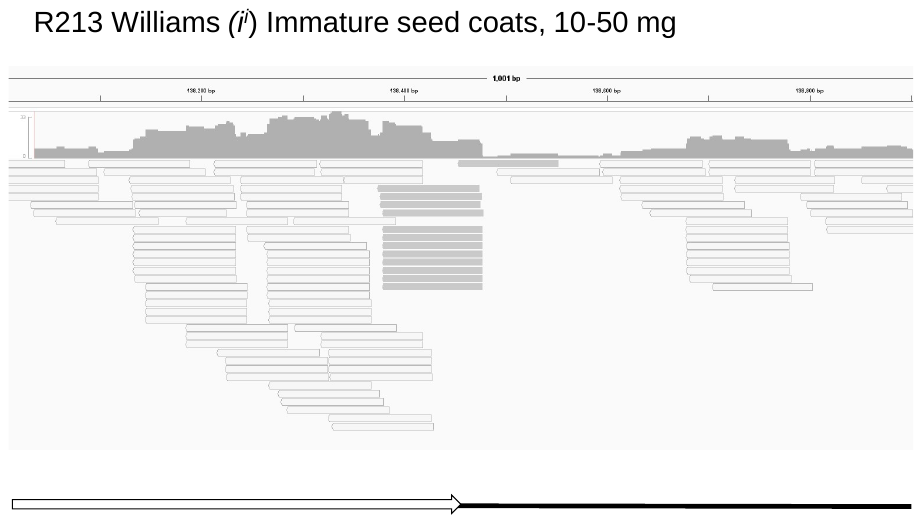

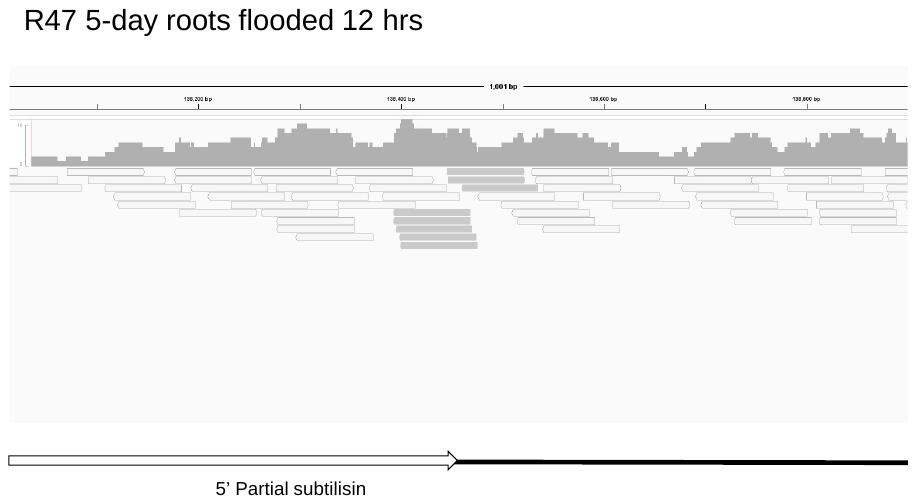

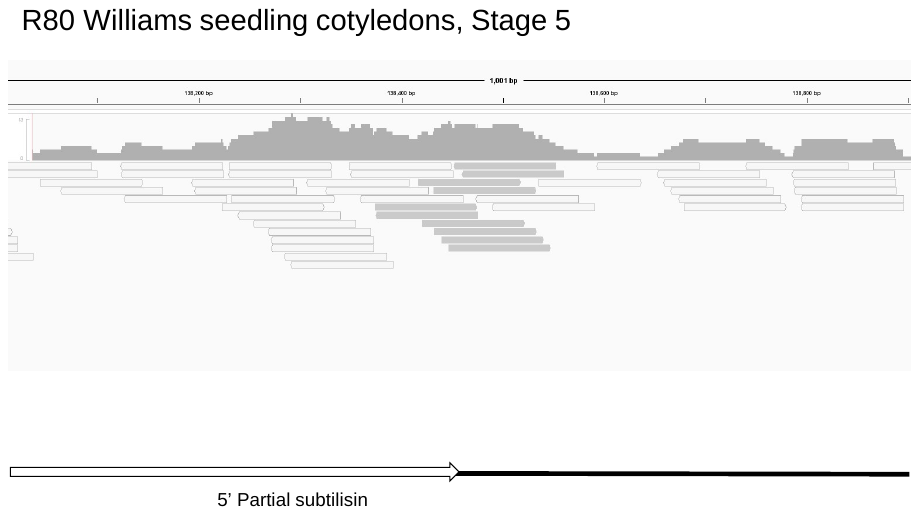

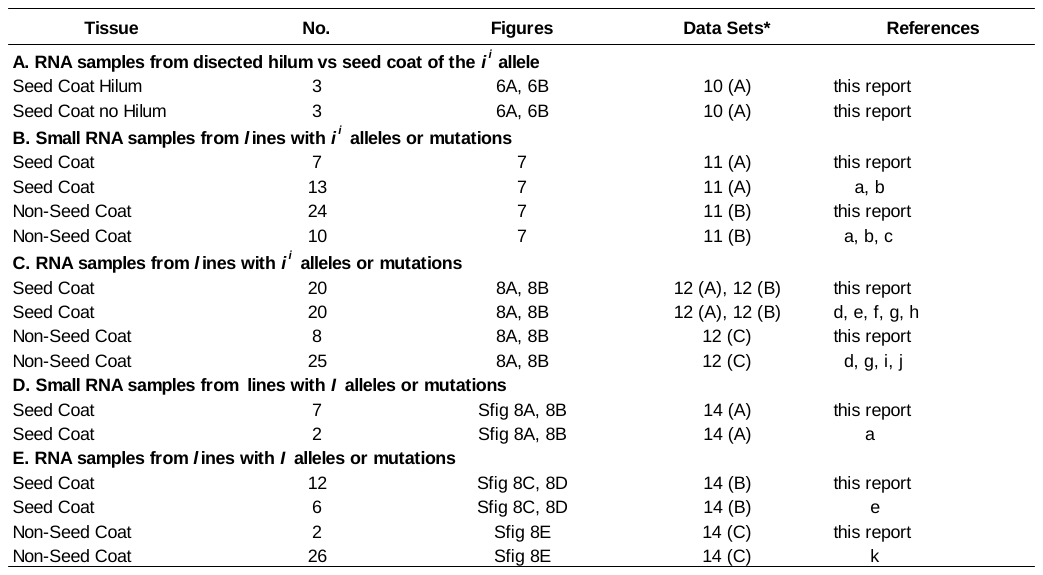

**Supplemental Table 4.** RNA-seq or small RNA-seq samples from seed coats versus non-seed coat tissues of I locus alleles. No. is the total number of samples subjected to high throughput sequencing, some of which are used in the indicated Figures,. The supplemental Data Sets contain the CHS siRNA levels or the transcript expression levels for the cognate subtilisin, DnaJ, and P450 Glyma model and all information on the samples. The data is first reported here of examined from the raw data reported in the following published references: (a)Tuteja et al., 2009; (b) Cho et al., 2013; (c) Zabala et al, 2012; (d) Jones et al., 2013; (e) Kour et al., 2014; (f) Zabala and Vodkin, 2014; (g) Cho et al., 2019; (h) Zabala et al., 2020; (i) Hunt et al., 2011; (j) Shamiuzzaman and Vodkin, 2014; (k) Jones et al., 2020.

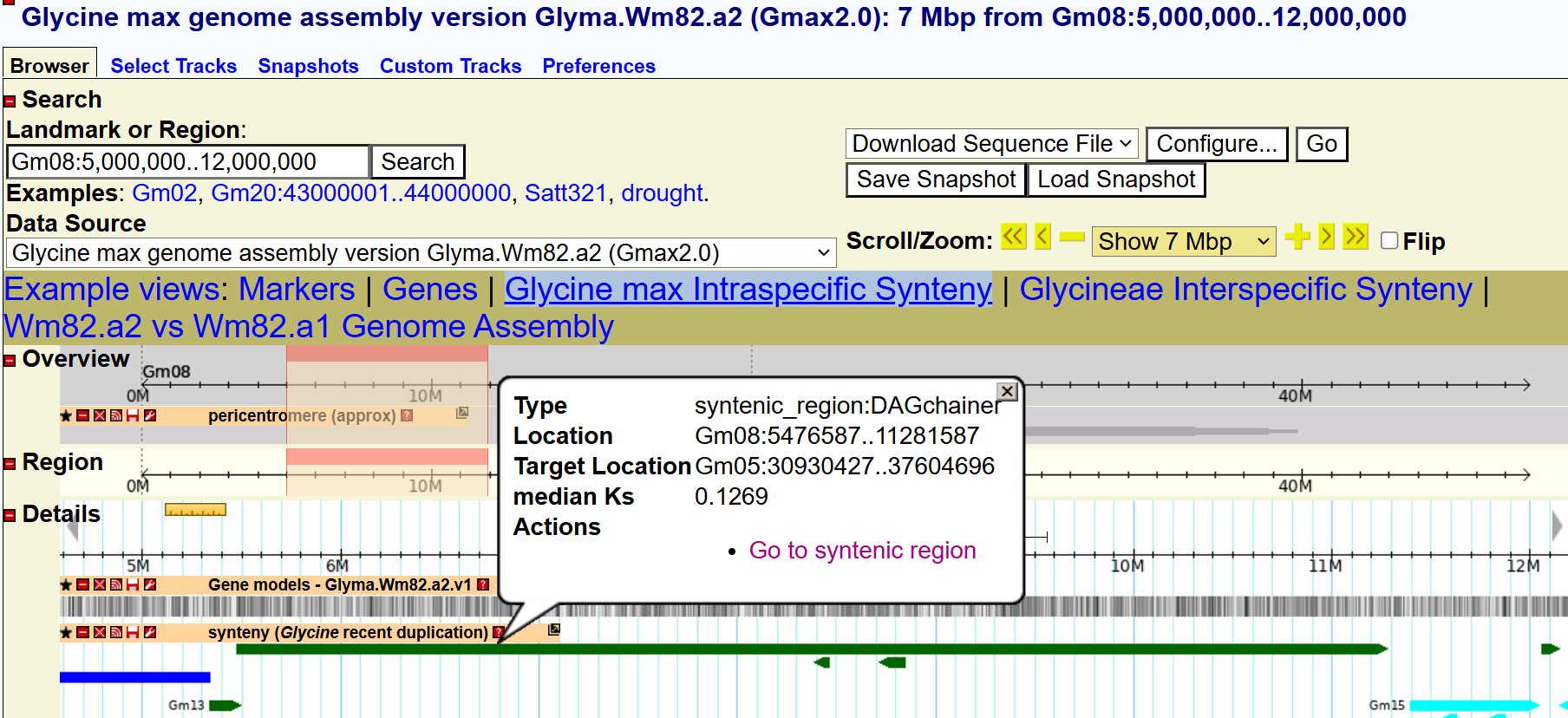

**Supplemental Figure 6.** Synteny analysis within soybean genomes reveals recent duplication events that gave rise to the *I locus* regions (*Gm08:5,476,587..11,281,587* and *Gm08:30,930,427..37,046,969*).
